# Deep Learning-coupled Proximity Proteomics to Deconvolve Kinase Signaling In Vivo

**DOI:** 10.1101/2025.04.27.650849

**Authors:** Kanchan Jha, Daichi Shonai, Aditya Parekh, Akiyoshi Uezu, Tomoyuki Fujiyama, Hikari Yamamoto, Pooja Parameswaran, Masashi Yanagisawa, Rohit Singh, Scott H. Soderling

## Abstract

Deconvolving the substrates of hundreds of kinases linked to phosphorylation networks driving cellular behavior is a fundamental, unresolved biological challenge, largely due to the poorly understood interplay of kinase selectivity and substrate proximity. We introduce KolossuS, a deep learning framework leveraging protein language models to decode kinase-substrate specificity. KolossuS achieves superior prediction accuracy and sensitivity across mammalian kinomes, enabling proteome-wide predictions and evolutionary insights. By integrating KolossuS with CRISPR-based proximity proteomics in vivo, we capture kinase-substrate recognition and spatial context, obviating prior limitations. We show this combined framework identifies kinase substrates associated with physiological states such as sleep, revealing both known and novel Sik3 substrates during sleep deprivation. This novel integrated computational-experimental approach promises to transform systematic investigations of kinase signaling in health and disease.

## Introduction

Deciphering the molecular logic of kinase-substrate relationships, connecting hundreds of kinases in mammalian proteomes to cellular behaviors through specific phosphorylation sites, is a central challenge in biology. Protein phosphorylation is one of the most evolutionarily conserved and prevalent post-translational modifications, with 30–70% of all proteins containing phosphorylated serine, threonine, or tyrosine residues (*1, 2*). Approximately 2% of the human genes are dedicated to encoding kinases, reflecting their critical roles in biology (*3*). Phosphorylation regulates nearly every biological phenomenon (*4–7*) including fundamental cell processes such as cell cycle (*8*), metabolism (*9–11*), gene expression (*12*), and high-level behaviors such as learning and memory (*13*). It achieves this by modulating protein activity, stability, localization, and interactions in response to cellular stimuli. Dysregulation of phosphorylation underlies a range of diseases, making kinases one of the most actively studied therapeutic targets (*14, 15*).

Kinase specificity is dictated by the twin factors of biophysics and subcellular localization. Recognition of a specific protein substrate is largely governed by local biophysical patterns that govern substrate recognition motifs around the phosphoacceptor site and kinase-substrate co-localization mechanisms. Substrate recognition motifs are determined by interactions with the catalytic domain of kinases, which consist of conserved β-sheet and α-helical structures that position substrates for phosphorylation (*16–18*), influenced by the biochemical properties of residues in this catalytic cleft. For instance, the PKA consensus sequence R-R-X-S/T-Φ (Φ = hydrophobic residue) fits precisely with side chain interactions within the kinase active site (*16, 19*). Recent large-scale studies using in vitro phosphorylation of degenerate peptide libraries or dephosphorylated lysates have begun to elucidate the substrate motif preferences of human kinases, but these studies are limited in their ability to generalize to kinases without data or to kinases across model organisms (*2, 20, 21*). Critically, they do not address the other essential determinant of in vivo kinase activity: the crucial role of spatial proximity between kinases and substrates that is required for kinase-substrate interactions in vivo (*22*). For example, substrates may possess perfect phosphoacceptor motifs for a given kinase, but if they are not within proximity to the kinase, they will not be phosphorylated.

Despite decades of research, 80% of kinases have fewer than 20 identified substrates, and many lack any known substrates entirely, categorized as “dark kinases” (*23*). Advances in phosphoproteomics have expanded the catalog of phosphorylation sites, yet the attribution of these modifications to specific kinases remains largely uncharted. Less than 5% of known human phosphopeptides have annotated kinases, limiting our understanding of the signaling networks that govern biological processes (*23*). Bridging these gaps requires novel approaches that integrate computational and experimental strategies to define kinase-substrate relationships comprehensively.

Deep learning has emerged as a promising tool for tackling this challenge, offering the ability to process large datasets and learn relevant data features and patterns autonomously. Traditional machine-learning approaches, relying on manually curated features such as structural or evolutionary properties, have shown limited success due to biases and incomplete feature sets. By contrast, deep-learning models can uncover rich representations from raw data, enabling significant advancements in kinase substrate prediction tasks. Early efforts, such as MusiteDeep (*24*), applied convolutional neural networks (CNNs) to predict phosphorylation sites, while later architectures like capsule networks (*25*) and densely connected CNNs have further improved prediction accuracy (*26*). However, these models’ performance degraded substantially on kinases outside their training data, as they failed to capture general substrate-specificity features across kinase families or species. More recently, large protein language models (PLMs) have revolutionized the field of protein biology by capturing the intricate dependencies within sequences using transformer-based architectures (*27–29*). These models encode protein sequences into contextual embeddings that capture structural and functional features, enabling applications such as protein-drug interaction prediction (*30, 31*), protein structure and function annotation (*32–35*), and post-translational site prediction (*36*). Early studies employing language models to predict kinase-substrate interactions demonstrate their potential (*36–39*), yet challenges remain in achieving generalizability across kinases and connecting these sequence-based predictions to the spatial organization of signaling networks.

Here, we present two advances to decode kinase function in vivo (Fig. 1A-B). 1) KolossuS, a deep-learning model that leverages embeddings from the evolutionary-scale protein language model ESM-2 to learn the sequence language of kinase-substrate selectivity from large datasets of in vitro serine, threonine, and tyrosine kinase assays. KolossuS achieves state-of-the-art performance in substrate prediction, capturing latent embeddings of kinase-specific motifs agnostic to species. 2) To resolve co-localization as a driver of kinase selectivity, we integrate KolossuS with proximity proteomics using a CRISPR-based in vivo BioID approach (*40*) coupled with phosphoproteomics. Applying this integrated framework, we found that kinases (CaMK1D and Brsk2), which display convergent substrate evolution, can share the same phosphorylation sites in vivo. Moreover, we demonstrate this approach can be coupled with behavioral manipulations in mice to deconvolve mechanisms downstream of the kinase Sik3 (*41*) relevant to sleep need.

**Fig. 1:**
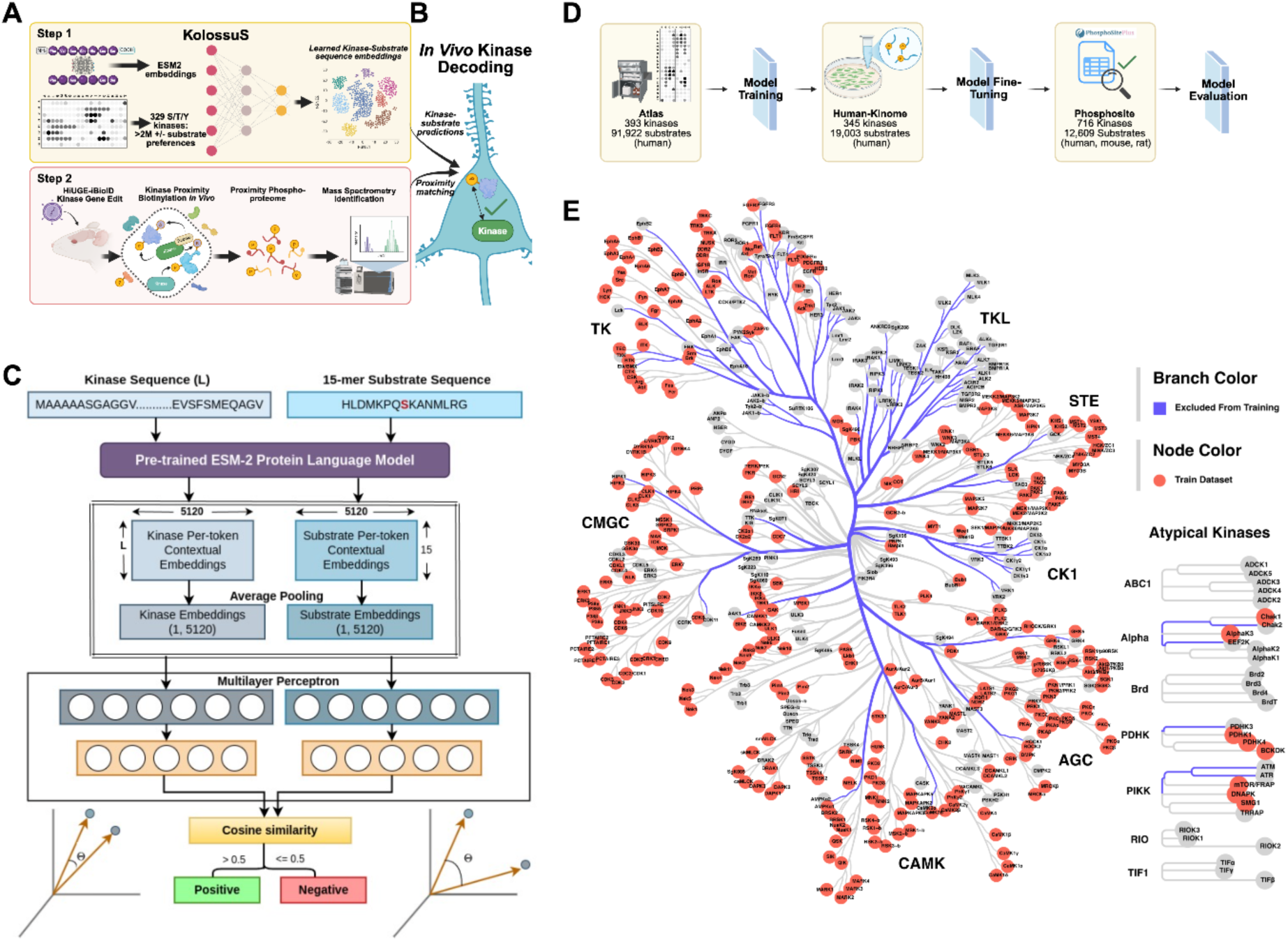
A Hybrid Computational-Experimental Framework for Mapping Kinase Substrate Networks Using KolossuS and In Vivo Proximity Labeling. **(A, *Step 1*)** KolossuS is a new deep learning model that leverages protein language models and experimental in vitro kinase-substrate data to learn kinase-substrate selectivity based just on protein sequences. (**A, *Step 2***) Kinase-substrate proximity in vivo is discovered using Homology-independent Universal Genome Engineering (HiUGE)-in vivo Biotin Identification (iBioID) to tag endogenous kinase with TurboID for biotinylation of nearby proteins. Biotinylated kinase proximal proteins are then digested and enriched for phosphophorylated peptides for phosphoproteomics. (**B**) KolossuS interprets the kinase-proximal phosphoproteomic data to reveal substrates. (**C**) Diagram of the KolossuS model architecture. Average-pooled kinase and substrate 15-mer embeddings are generated from a pre-trained ESM-2 PLM and non-linearly projected by the multilayer perceptron into a shared co-embedding space, in which the probability of the kinase phosphorylating the substrate is captured by the cosine similarity between their representations. (**D**) Schematic of KolossuS training and benchmarking. KolossuS is trained on broad-scale in-vitro PSPA data to capture broad patterns of kinase-substrate specificity (Atlas), and fine-tuned on LC/MS data capturing kinase-substrate interactions in a cellular context (Human Kinome). The final evaluation is performed on a dataset containing literature-curated phosphorylation events withheld from the other two datasets (Phosphosite). (**E**) Phylogenetic tree of kinases included within the training (Atlas) datasets. KolossuS is trained using phylogenetic-aware splits in the training and validation/testing datasets. Kinases seen during training are indicated using red; blue edges indicate kinases held out from the training dataset. Note that the TKL and CK1 families are excluded entirely from both the training and fine-tuning dataset. The structure of the phylogenetic tree was obtained from (http://phanstiel-lab.med.unc.edu/CORAL/).

This work opens new windows into the biology of phosphorylation signaling within living animals by holistically addressing two important barriers—kinase selectivity and substrate proximity—to untangling kinase biology in vivo. To facilitate broader use, KolossuS is accessible via a web-based platform at https://singhlab.net/kinase

## Results

### KolossuS Model Overview

Predicting which proteins a kinase will phosphorylate remains a fundamental challenge in cell signaling research. Existing approaches to model kinase specificity suffer from several critical limitations: they often don’t generalize across the entire kinome (e.g., methods limited to just Ser/Thr kinases (*20*), fail to capture complex interactions between substrate residues (e.g., position weight matrices that evaluate each location independently (*20, 21, 42*), and most crucially, lack proper calibration across different kinases (*43*). The calibration problem significantly hampers practical applications—a model might correctly rank substrate preferences for individual kinases yet assign inaccurate absolute scores across kinases, making it impossible to reliably match phosphoproteomics data to their cognate kinases. Consider the following toy example: suppose substrate S_1_ is phosphorylated in vivo by kinase K_1_, and S_2_ by K_2_. Imagine a poorly calibrated model that predicts the following probabilities: p(K_1_,S_1_)=0.2, p(K_1_,S_2_)=0.1, p(K_2_,S_1_)=0.8, p(K_2_,S_2_)=0.9. For each kinase individually, the model is accurate in ranking substrate preferences. Yet, given a phosphoproteome where S_1_ and S_2_ were observed, it will incorrectly associate them both to K_2_. KolossuS addresses all of these limitations while also interpreting the underlying organization of kinase specificity across evolutionary distances and substrate preferences.

KolossuS harnesses the power of protein language models to understand the sequence grammar of kinase-substrate recognition. Trained on millions of protein sequences, these models capture evolutionary patterns that encode key structural and functional relationships, proving effective for protein design, interaction prediction, drug binding, and allosteric regulation (*30, 43–49*). We use these models to generate initial representations of kinases and substrates, which we then iteratively fine-tune using both in vitro and in vivo datasets. Our approach emphasizes interpretability and generalizability while maximizing the capabilities of the underlying language models. Specifically, we implement a co-embedding architecture that represents kinases and substrates in an interpretable, shared mathematical space—a strategy that enables both accurate kinase-substrate specificity predictions and reasoning about evolutionary relationships across the kinome.

KolossuS operates with sequence-only inputs, making it directly applicable to phosphoproteomics assays (Fig. 1C). The model processes the full kinase sequence while representing substrates as 15-mer peptides centered at the phosphorylation site. This substrate representation strategy reflects several key considerations: phosphorylation is primarily governed by the local physical and electrostatic context surrounding the site (*50*); recent atlas-scale assays (*2, 20, 21*) provide abundant k-mer level data; and phosphoproteomics typically reports peptide fragments rather than full proteins. We extract initial representations using the ESM-2 15B parameter model, aggregate per-residue embeddings into consolidated kinase and substrate representations, and nonlinearly transform these into a shared embedding space. During training, the model learns to position interacting kinases and substrates close to each other in this latent space. This approach encodes the hypothesis that kinases with similar representations will phosphorylate similar substrates, creating an interpretable framework that captures functional relationships across the kinome.

### KolossuS predicts kinase-substrate interactions with unprecedented accuracy and generalizability

The design of KolossuS reflects the multi-scale ecosystem of kinase-substrate data, spanning from atlas-scale synthetic sequence assays to focused cellular phosphorylation studies. We reserved literature-based substrate data for benchmarking while using broader, in vitro datasets for training and fine-tuning (Fig. 1D). First, we trained on “Atlas” datasets generated by Johsnon et al. and Yaron-Barir et al. (*20, 21*), containing predicted interactions for 393 human kinases derived from position-specific scoring matrices (PSSMs) from synthetic peptide array experiments. These datasets provided extensive coverage but are based on data from non-natural sequences. We then fine-tuned our model using Sugiyama et al.’s “Human Kinome” dataset (*2*), which contains experimentally determined substrates for 354 human kinases identified through LC/MS-based in vitro assays of cellular proteins. This progressive training approach allowed us to capture both the breadth of potential kinase-substrate interactions and the precision of experimentally determined natural sequence sites.

To rigorously evaluate KolossuS’s generalizability, we implemented phylogeny-aware data splits rather than simple sequence similarity-based partitioning (Fig. 1E; Materials and Methods). We designated an approximately 80:10:10 train:validation:test split, with the CK1 and JAK kinase families reserved for validation and the TKL and FAK families held exclusively for testing. The CK1 and TKL families are Ser/Thr (S/T) kinases while the other two families are Tyr (Y) kinases. Additionally, we withheld select kinases from each family in the training set, allocating them to validation or testing only. This stratification enabled us to evaluate not only KolossuS’s ability to interpolate within kinase families but also extrapolate to evolutionarily distinct kinases. For final benchmarking after fine-tuning, we used data from Phosphosite (*51*), which contains experimentally validated kinase-substrate interactions from human, mouse, and rat literature.

KolossuS demonstrates state-of-the-art performance on kinase-substrate prediction, achieving an AUROC (area under receiver operating characteristic) of 75.5% (83.8%) for Ser/Thr (Tyr) human kinases (Fig. 2A). KolossuS remains comparably accurate on mouse and rat phosphorylation (Fig. 2B,C). The two-stage training process is also effective: on the human kinases present in our evaluation dataset, zero-shot prediction with average-pooled ESM-2 embeddings achieves an AUROC of 50.3% (i.e., essentially random) and AUPR (area under precision-recall curve) of 42.5%. Supervising with just the atlas data (“pre-fine-tuned”) improves these metrics to 75.4% and 66.7%, respectively (Fig. 2D, Table S1). Fine-tuning on the Human Kinome data further improves AUROC (AUPR) to 76.2% (67.9%). We also compared KolossuS against Phosformer-ST (*37*), currently the best-performing kinase-specificity prediction method (Fig. 2E, Table S2). Phosformer-ST also leverages PLMs and is trained on atlas data but is limited to S/T kinases. On these kinases, KolossuS achieves higher accuracy, sensitivity, AUROC and AUPR. Its outperformance over Phosformer-ST is stronger in the case of Tyr kinases, which the latter was not trained on. Even on held-out kinase families not seen during training, KolossuS demonstrates robust performance, consistently outperforming Phosformer-ST despite the expected degradation in accuracy (Fig. S1).

**Fig. 2:**
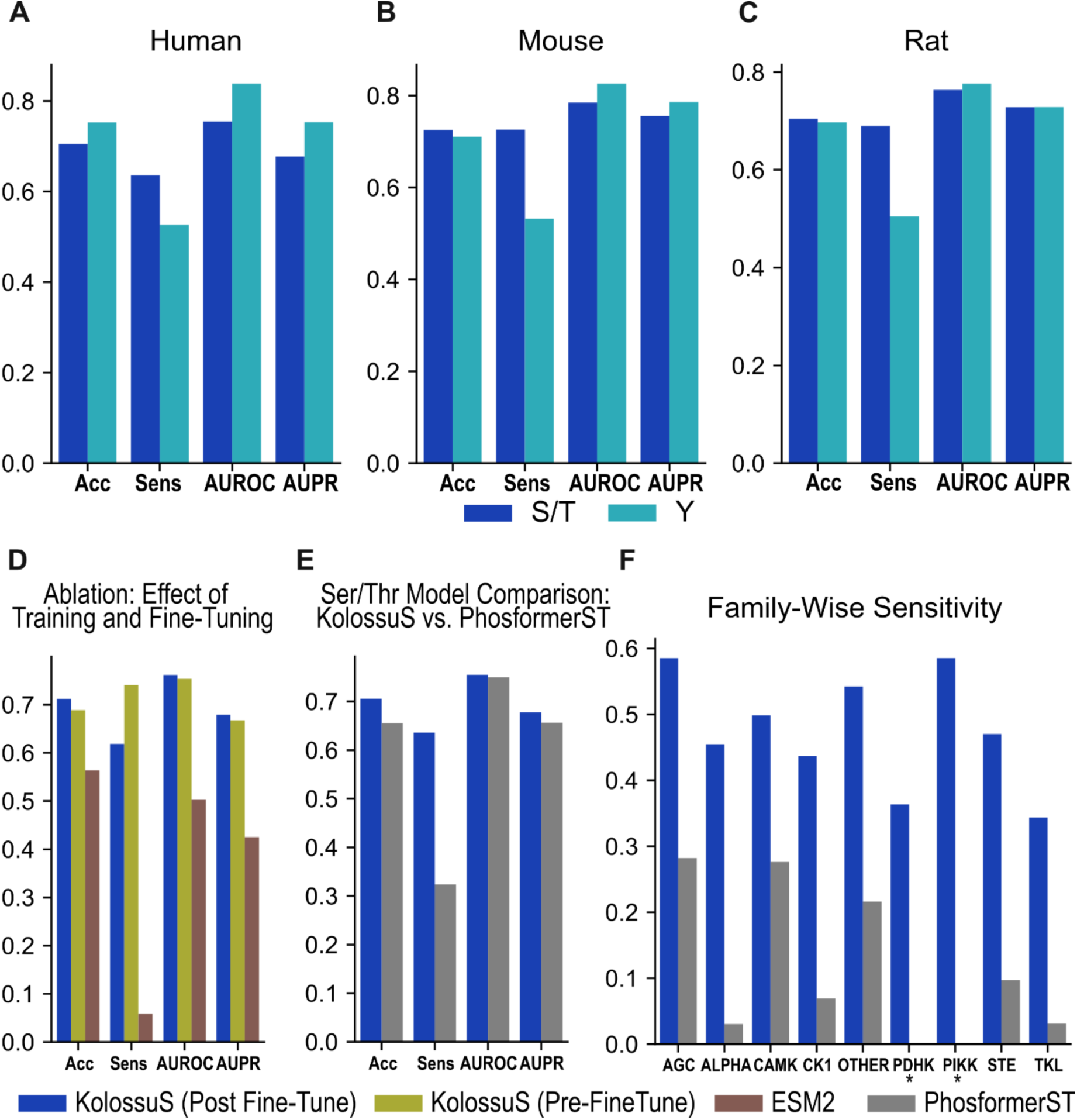
Summary of KolossuS performance on the Phosphosite evaluation dataset. **(A-C)** KolossuS performance on Ser/Thr (S/T) and Tyr (Y) kinases in human, mouse and rat, respectively. Acc: Accuracy; Sens: Sensitivity; AUROC: Area under the receiver operator characteristic curve; AUPR: Area under the precision-recall curve. **(D)** Comparison of prediction performance between KolossuS (pre- and post-fine tune) and raw ESM-2 embeddings on the human kinases in the Phosphosite evaluation dataset (mouse and rat data available in Table S1). **(E)** Performance comparison between KolossuS and Phosformer-ST on the S/T human kinases (mouse and rat data: Table S2). **(F)** Comparison of model sensitivity between KolossuS and Phosformer-ST across 9 Ser/Thr families (human only). Families were chosen such that data was available for at least 5 human kinases and 40 kinase-substrate pairs in the Phosphosite dataset. Asterisks indicate families for which Phosformer-ST had a sensitivity of zero (no correctly identified substrates).

One of KolossuS’s key advantages lies in its superior calibration—the alignment between predicted probabilities and actual outcome frequencies. This calibration translates directly to practical benefits in laboratory settings, enabling the matching of phosphoproteomes to cognate kinases. KolossuS is substantially more sensitive than Phosformer-ST when evaluated at the same threshold of p = 0.5: the latter identifies fewer than 50% of true positive S/T substrates across human, mouse, and rat datasets, with uneven performance across specific kinase families. For example, Phosformer-ST fails to identify any positive pairs for the PDHK and PIKK families, while KolossuS correctly predicts 59% of positive pairs (Fig. 2F). To systematically assess calibration quality, we calculated the optimal prediction threshold for each family. Specifically, we computed the F1-max threshold, the prediction threshold at which the F1 score (harmonic mean of sensitivity and specificity) reaches its maximum. A perfectly calibrated model will achieve F1-max at 0.5 as the prediction threshold. For Phosformer-ST, family-specific F1-max scores are achieved at thresholds near 0.0. In contrast, KolossuS reaches optimal performance at thresholds between 0.15-0.37 (Table S3), substantially closer to the ideal 0.5 threshold of perfect calibration.

Ablation studies confirm the robustness of KolossuS’s architecture and training approach. We assessed base PLM models of different parameter sizes, finding that the ESM-2 15B parameter model offered the best performance. The smaller ESM-2 variant, with 650M parameters, performed slightly worse. Interestingly, a newer but slightly smaller model, ESM-C 6B (*52*), also slightly underperformed the older 15B model. As the lower computational cost of the 650M model may be valuable in GPU-constrained settings (Fig. S2), we make trained model weights available for both the ESM-2 650M and 15B variants.

### KolossuS embeddings reveal functional convergence and divergence across kinase families

KolossuS enables the first comprehensive prediction of the human kinome-substrate landscape, providing unprecedented insights into phosphorylation networks at genome scale. We generate de-novo predictions between 466 known active human kinases (458 mouse kinases) and 227,736 human substrate sequences (100,981 mouse substrate sequences). The latter represent the entirety of substrates reported in Phosphosite+ (Materials and Methods). Using a prediction score threshold of 0.75 to designate high-confidence positive predictions, we predicted phosphoproteomes for both human and mouse. These predictions suggest most kinase-substrate interactions are highly specific: the vast majority of substrate sequences (>90%) are targeted by fewer than 11% of kinases (Fig. S3). This pattern of selective recognition holds consistently across both species, suggesting evolutionary conservation of kinase-substrate specificity patterns.

The design of KolossuS encodes a biological hypothesis: kinases and substrates can be represented in a shared latent space where vector distance (measured by cosine similarity) indicates compatibility. This allows us to construct a unified mathematical framework for reasoning about substrate recognition and kinase functional relationships. While representing complex biochemical interactions through vector similarity is a simplifying assumption, the expressivity of PLMs allows us to learn a rich latent space which captures meaningful biological patterns and uncover functional relationships that transcend traditional phylogenetic classifications. We visualize these relationships through complementary approaches: UMAP projections reveal global patterns in kinase organization, while dendrograms that allow us to compare substrate specificity patterns with evolutionary relationships (*3*). Analyzing these visualizations reveals previously unrecognized patterns of functional organization. For instance, we observe a clear separation between the CMGC family of kinases and other S/T kinase families in our UMAP visualization (Fig. 3A). This separation is absent in the ESM-2 embedding space (Fig. 3B), suggesting it arises specifically from substrate preferences rather than structural properties. This finding aligns with the distinctive substrate profile of CMGC kinases, which predominantly target +1 proline or arginine-directed phosphorylation sites, are evolutionarily ancient, and contain a unique catalytic domain insert that may regulate kinase activation (*43, 53*). Overall, KolossuS-derived embeddings better agree with phylogenetic groupings than ESM-derived groupings, broadly supporting the phylogenetic classification of kinase function. However, we observed that the AGC and STE families show increased divergence in KolossuS embeddings, suggesting divergence in substrate specificities within those families (Fig. 3C; Fig. S4; Discussion).

**Fig. 3:**
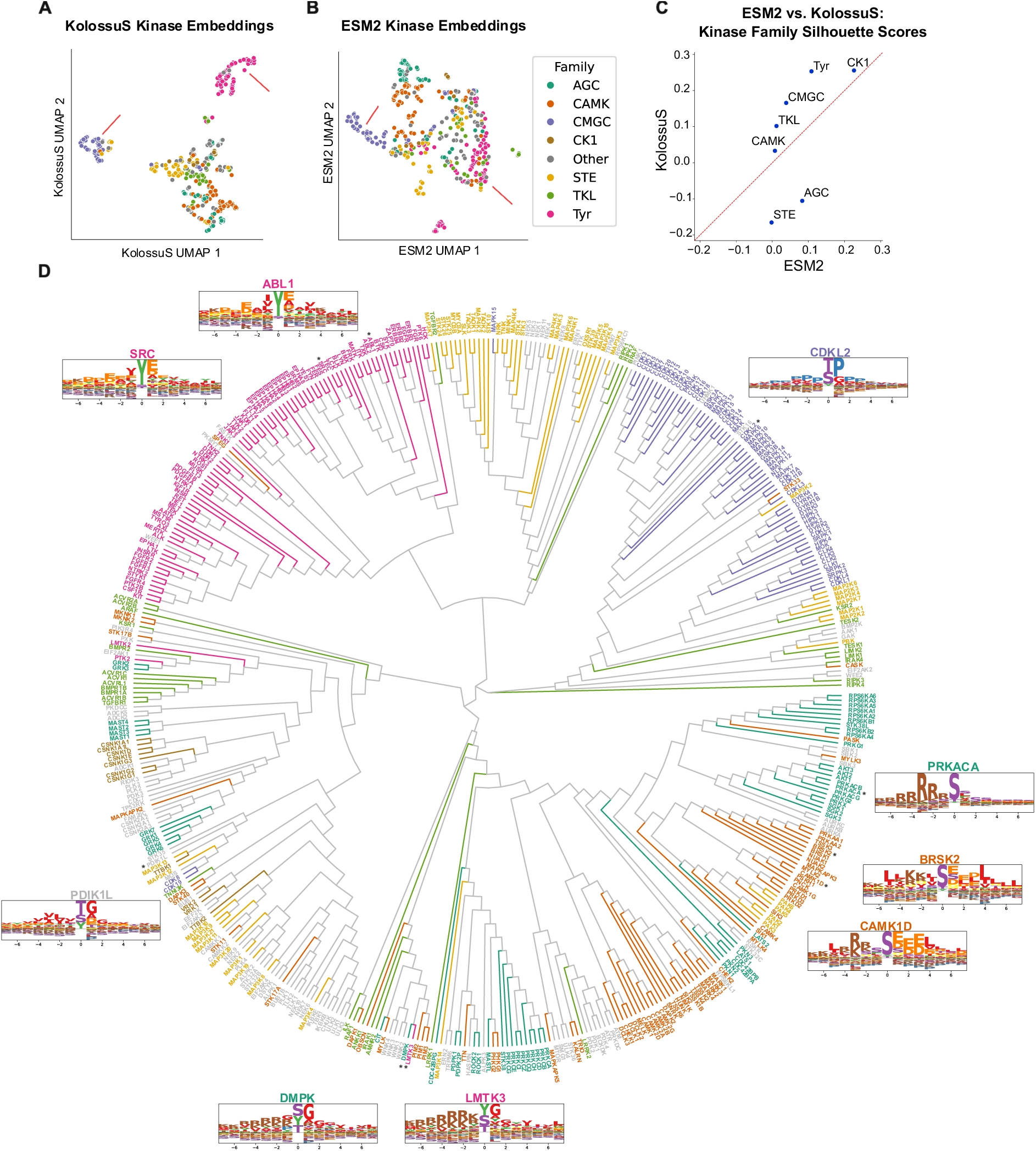
Visualization of KolossuS embeddings and substrate binding motifs. **(A-B)** UMAP visualization of 466 known human kinases using KolossuS and ESM-2 embeddings, respectively. Red arrows: KolossuS embeddings indicate clear separation between the CMGC from the other S/T kinase families, and a clear separation between Tyrosine (Tyr) kinases. **(C)** Scatterplot of silhouette scores calculated for clustering of kinases according to family in the KolossuS and ESM-2 embedding spaces. The red diagonal line indicates an equal silhouette score between the two embedding spaces. **(D)** Dendrogram of kinases created from the KolossuS kinase embeddings, colored by kinase families (same as in panel **A**). A distance matrix was created using the cosine similarity metric, and a hierarchical clustering was computed using the neighbor-joining method (Materials and Methods). Predicted motifs are displayed for select kinases (indicated with asterisks in the dendrogram): the PKA (PRKACA) predicted motif is consistent with its established motif; CDKL2 displays the proline-directed phosphorylation motif characteristic of CMGC kinases; PDIK1L is a dark kinase with no known motif; SRC and ABL1 are two well-studied tyrosine kinases involved in cancer, our predicted motifs for these kinases are consistent with prior studies (Fig. S8). Colors on logos represent the amino acid chemical properties: hydrophobic (red), aromatic (green), polar (purple), acidic (orange), amide (yellow), basic (brown), Proline (blue).

The KolossuS-derived dendrogram also shows stronger family-based clustering than an equivalent dendrogram based on ESM-2 embeddings (Fig. 3D, Fig. S5). This indicates that the traditional phylogenetic classification of kinases may indeed capture functional substrate preferences more accurately than whole-protein representations (which ESM-2 proxies). However, it again shows greater dispersion of STE and AGC kinase embeddings across the dendrogram, re-indicating divergence within those families.

KolossuS also reveals instances of convergent evolution across the kinome. Members of distinct kinase families sometimes cluster together in our embedding space, indicating they have evolved similar substrate recognition patterns. An example is the close pairing of DMPK (an S/T kinase from the AGC family) and LMTK3 (a tyrosine kinase) in our dendrogram (Fig. 3D). Despite their distinct catalytic domains and evolutionary origins, both kinases appear to recognize remarkably similar substrates. Gene Ontology enrichment analysis of their shared predicted substrates revealed significant overrepresentation of cytoskeletal and actin regulatory processes, including components of the Rho GTPase signaling pathway (Fig. S6). This functional convergence aligns with their established biological roles—LMTK3 regulates cell migration through cytoskeletal dynamics in cancer contexts (*54*), while DMPK modulates muscle contractility through actin-myosin signaling (*55*). We note that such substrate preference similarity doesn’t necessarily imply identical biological function, as subcellular localization and expression patterns also critically determine a kinase’s actual physiological targets.

Unlike classical prediction approaches that model each substrate residue position independently, KolossuS can capture complex dependencies between residue positions. Even though we average-pool substrate embeddings, the representational power of the underlying PLM encodes crucial positional information in the average, as confirmed by a position-specific entropy analysis (Fig. S7). KolossuS enables us to extract interpretable motif representations of kinase specificity. We generated sequence logos for each kinase from our predicted phosphoproteome, visualizing the frequency and positional preference of amino acids surrounding phosphorylation sites. The resulting motif logos for well-characterized kinases like PKA (Fig. 3D) closely match their established recognition patterns. Additionally, our predicted motifs correctly identify the established proline-directed phosphorylation site that characterizes CMGC binding motifs. Similarly, the predicted motifs for well-studied Tyrosine kinases, e.g. SRC and ABL1, are also consistent with the literature (Fig. S8)(*56–58*). Crucially, KolossuS newly enables motif determination for understudied “dark kinases” lacking experimental characterization, providing the first comprehensive atlas of predicted substrate motifs across the entire human kinome. We make these predictions available upon request.

Kinases located near each other in the dendrogram exhibited similar motif logos. For example, Brsk2 (Brain-specific kinase 2, also known as SAD-B) and CaMK1D (Calcium/Calmodulin Kinase 1-delta) clustered within neighboring clades of the dendrogram and displayed similar characteristics in their predicted motifs (Fig. 3D): a preference for non-polar amino acid residues at the -5 position, basic residues (Arginine and Lysine) at the -3 and -2 positions, and acidic residues (Aspartic Acid and Glutamic Acid) at positions +1 through +3. Both kinases belong to the large CAMK kinase family; however, Brsk2 does not bind calmodulin and is part of the AMPK-related subfamily, whereas CaMK1D is activated by calcium-calmodulin and its upstream kinase, CaMKK, and belongs to the CAMK1 subfamily.

Brsk2 is evolutionarily conserved and was identified in C. elegans (termed Sad-1), where it regulates neuronal polarity and presynaptic vesicle clustering (*59*). In mice, Brsk2 phosphorylates the axonal protein Tau as well as the presynaptic protein Rim1, and its loss affects neuronal polarity during development (*60–62*). Consistent with its developmental and synaptic roles, genetic variants of Brsk2 are associated with developmental delay, autism, and intellectual disability in humans (*63, 64*).

In neurons, CaMK1D is localized across several subcellular compartments, including synapses and the nucleus, where it phosphorylates a broad range of substrates (*65*). At synapses, CaMK1D binds the scaffolding protein Git1, facilitating its phosphorylation of the substrate ArhGef7 (βPIX), which modulates actin cytoskeletal dynamics via Rac1 during synaptogenesis (*66*). CaMK1D phosphorylates Ras-GRF1, activating ERK signaling during long-term potentiation (*67*). CaMK1D also phosphorylates eIF4G, modulating protein translation (*68*). Although there is no direct evidence linking Brsk2 and CaMK1D, CaMK1D similarly regulates aspects of early neuronal development and polarization, analogous to Brsk2 (*69*). Interestingly, studies of CaMKK have shown that, like CaMK1D, Brsk2 can also be phosphorylated in its activation loop, leading to CaMKK-mediated stimulation of Brsk2 kinase activity (*70*). This suggests that CaMK1D and Brsk2 may regulate similar signaling pathways and possibly substrates.

### *In vivo* analysis of kinase substrates using HiUGE-iBioID and KolossuS

To experimentally test the possibility that CaMK1D and Brsk2 share overlapping functions/substrates in vivo, we employed Homology-independent Universal Genome Editing iBioID (HiUGE-iBioID), a recently developed proximity-labeling method combining CRISPR-Cas9 gene editing and TurboID-based biotinylation in living mice. (Fig. 4A) (*40, 71*). Specifically, we investigated five kinases (Brsk2, CaMK1D, as well as Aak1, Akt1, and Uhmk1). These were selected based on their diverse family classifications and robust expression within brain tissues. We reasoned the resulting kinase-TurboID fusion proteins would mediate proximity-dependent biotinylation of nearby proteins, including their substrates, enabling their identification from vivo. If successful, this would address the critical contribution of proximity to kinase-substrate interactions.

**Fig. 4.**
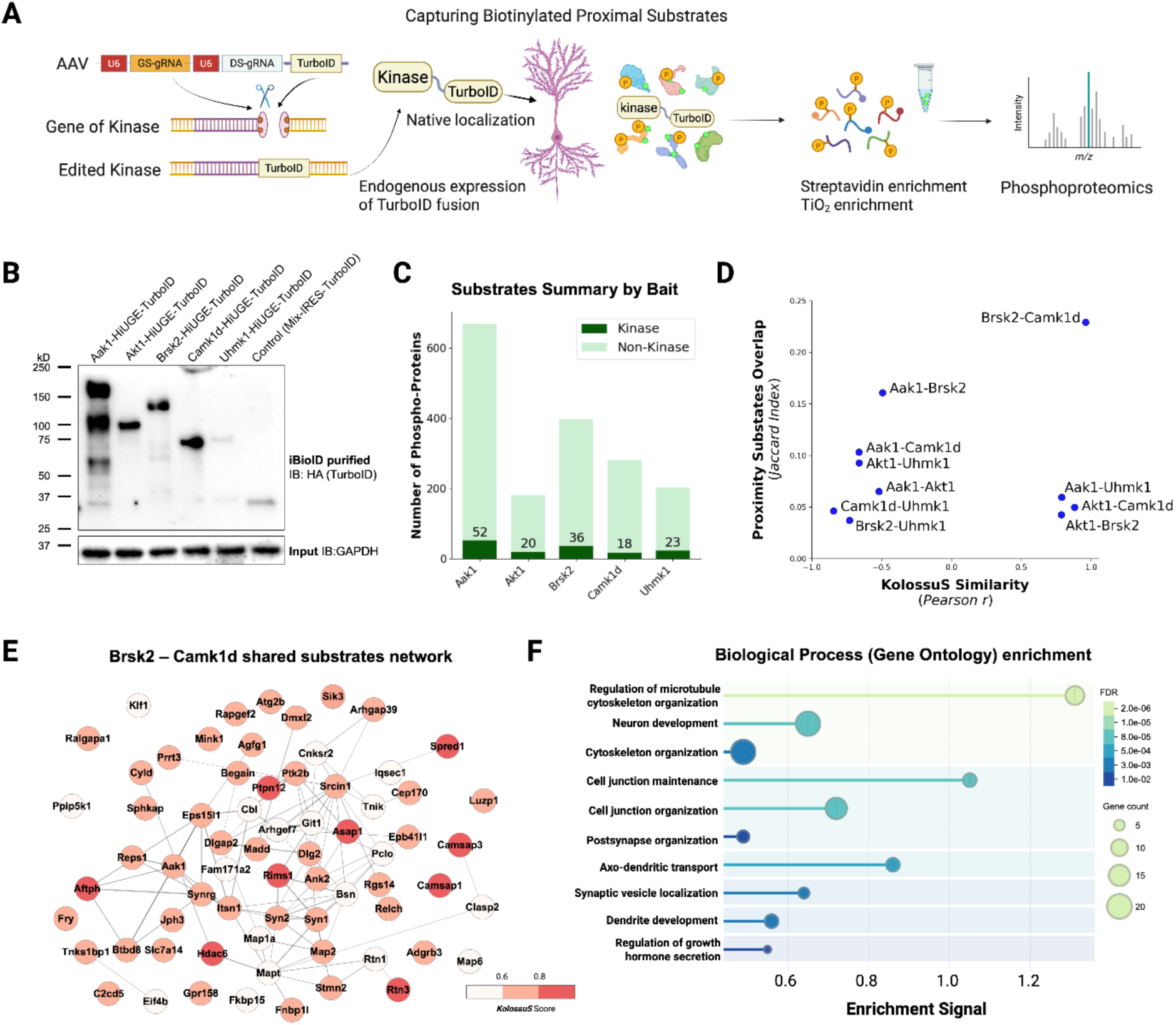
Combining HiUGE-iBioID with KolossuS Predictive Scoring Identifies Direct Kinase Substrate Networks In Vivo. **(A)** Schematic illustration of the AAV-mediated HiUGE-iBioID workflow, followed by streptavidin pulldown and TiO₂ enrichment for proximity phosphoproteomics. **(B)** Western blot showing endogenous tagging of TurboID-HA via HiUGE for various kinases in mouse brain tissue, alongside a soluble TurboID-HA control (HiUGE-IRES-TurboID-HA). **(C)** Bar graph summarizing the number of proximal phosphoproteins identified for each bait kinase, with counts of kinase and non-kinase substrates. Proximal phosphoproteins were defined as phosphopeptides enriched in TurboID-fusion samples relative to the soluble TurboID control (log₂FC > 2, Bonferroni-adjusted p < 0.05). **(D)** Scatterplot comparing KolossuS similarity (x-axis; Pearson correlation of predicted scores for shared phosphopeptides by a given pair of kinases) and phospho-substrate overlap (y-axis; Jaccard similarity index) across all kinase pairs. **(E)** Network of high-confidence shared substrates between Brsk2 and CaMK1D (KolossuS score > 0.50 for both). Node color indicates the average KolossuS score from both kinases; edges represent STRING interactions at medium confidence (0.40). **(F)** Gene Ontology (GO) Biological Process enrichment of Brsk2-CaMK1D shared substrates (KolossuS > 0.50 in both), with enriched terms clustered by Jaccard similarity using the STRING API (https://string-db.org/).

Postnatal day 0–2 pups transgenic for Cas9 expression received bilateral intracranial injections of purified PHP.eB-packaged AAV expressing gRNAs specific for each kinase and to the vector to liberate TurboID for NHEJ insertion into each locus. After 8 weeks of expression, mice were treated with biotin for five days. Controls expressed soluble TurboID, ensuring the specificity of identified proteins to each kinase (Fig. 4B). Following proximity labeling, biotinylated proteins were isolated using streptavidin affinity purification, and phosphorylated peptides from kinase proximal proteins were enriched using TiO₂ chromatography.

LC-MS/MS analysis identified kinase-specific phosphopeptides defined by a fold-change ≥1.5 and p-value <0.05 (two-tailed heteroscedastic t-test) relative to controls. This analysis yielded 1139, 211, 1023, 641, 271 unique phosphopeptides derived from 674, 182, 400, 284, 205 unique proteins for Aak1, Akt1, Brsk2, CaMK1D, and Uhmk1, respectively (Fig. 4C, Table S4). Notably, 21.85% of the identified phosphopeptides were not listed in the PhosphositePlus database, underscoring the potential of HiUGE-iBioID focused to a kinase to detect phosphorylation events that conventional phosphoproteomics approaches typically miss. The phosphopeptide profiles of Brsk2 and CaMK1D also displayed a high degree of similarity, compared to the other kinases (Fig. 4D). Of the peptides identified, 297 were detected in proximity to both Brsk2 and CaMK1D. Further analysis of the shared substrates between Brsk2 and CaMK1D demonstrated a strikingly high correlation (Pearson’s r = 0.960) between KolossuS prediction scores for Brsk2 and CaMK1D, strongly suggesting convergent substrate preferences (Fig. 4D,E).

While HiUGE-iBioID successfully identified a broad network of potential kinase-substrate interactions, precise mapping of kinase-substrate relationships remained challenging due to the presence of multiple kinases in proximity. Consistent with the notion that kinases can be clustered spatially via anchoring/scaffold proteins, lipids, and organelles; often in cascades, 80 out of the 960 quantified proteins (8.3%) were themselves kinases, complicating the identification of direct substrates of each bait kinase. To resolve this complexity, we applied KolossuS predictions to narrow down candidate kinase-substrate pairs. The combined HiUGE-iBioID and KolossuS analysis identified many known substrates of the target kinases, including Rim1 and Tau for Brsk2 and ArhGef7, Git1, Hdac5, eIF4G, and Ras-Grf1 for CaMK1D (Fig. S9, S10). Between Brsk2 and CaMK1D, KolossuS predictions identified 96 substrates (KolossuS score ≥0.50) common to both kinases (Fig. 4E). Gene ontology enrichment analysis of these dual-target substrates underscored the likely shared biology of these kinases with regards to neuronal development, microtubule regulation, and both pre- and postsynapse function (Fig. 4F). Several of these terms fit with prior literature for each kinase separately.

Kinase substrates can be boutique with respect to species, cell-type, and state and thus differ between species and cell types across studies (*72–75*). Our analysis of substrates specific to brain cell types revealed insights regarding these substrates for the remaining kinases, with several instances of known substrates or substrates aligned with their proposed cellular functions. *Uhmk1* (also termed KIS) is a novel RNA splicing regulatory kinase (*76*). HiUGE-iBioID-KolossuS detected likely phosphorylation of Ccar2, a core component of the DBIRD complex, a multiprotein complex that coordinates transcript elongation and alternative splicing (*77*). Other RNA binding and regulatory proteins included Hnrnpd (*78*), Cpeb4 (*79*), Atxn2 (*80*), and Ankrd17 (*81*). Of note, five substrates detected in brain by HiUGE-iBioID-KolossuS were also detected as Uhmk1 substrates in NIH3T3 cells by loss and gain of function phosphoproteomics (Scrib, Add1, Palm, Eef1d, and Ankrd17)(Fig. S11)(*76*). *AAK1* (adaptor-associated kinase 1) regulates clathrin-mediated endocytosis (*82*). Consistent with this function, Amph (amphyphysin) and NSF (N-ethylmaleimide-sensitive factor), which regulate clathrin endocytosis and synaptic vesicle recycling were detected and predicted as substrates. (Fig. S12) *AKT* (also, PKB) is known to phosphorylate beta-catenin (*83*), delta-catenin (*84*), and PAK1 (*85*), all among the substrates detected by HiUGE-iBioID and selected by KolossuS. (Fig. S13)

Together, these results support the notion that HiUGE-iBioID can capture substrates proximal to kinases in vivo. When paired with analysis by KolossuS, this combined approach appears to effectively address both major drivers of kinase-substrate interactions *in vivo*: kinase proximity and substrate motif matching.

### HiUGE-iBioID and KolossuS reveal *in vivo* substrates of Sik3 triggered by sleep deprivation

We next wondered whether this combined approach could be paired with in vivo substrate analysis under different physiological contexts that alter cell state and kinase activity. To test the feasibility of this, we analyzed the substrates of the kinase Salt-Inducible Kinase 3 (Sik3). Sik3 plays a pivotal role in regulating sleep homeostasis by modulating phosphorylation events influencing sleep need and non-rapid eye movement sleep (NREMS)(*41, 86*). Sleep deprivation induces Sik3 activity, mirrored by the hyperphosphorylation patterns observed in the Sik3 gain-of-function mutant (*Sleepy* mice) and implicating Sik3 as a critical regulator in sleep-related phosphorylation networks (*87*). However, the specific substrates and phosphorylation networks directly regulated by Sik3 during sleep deprivation remain under active investigation.

Similar to above, Cas9 knock-in mice received bilateral intracranial injections of AAV HiUGE-iBioID constructs targeting endogenous Sik3 for TurboID fusion. Eight weeks later, mice were administered biotin and subsequently subjected to 6 hours of sleep deprivation at the onset of the light phase (ZT0) to activate endogenous Sik3 or allowed ad libitum sleep conditions (Fig. 5A). Negative controls received identical sleep deprivation and biotin treatment, but expressed a soluble TurboID without Sik3 fusion, ensuring specificity of identified proteins to Sik3 proximity (Fig. 5B).

**Fig. 5.**
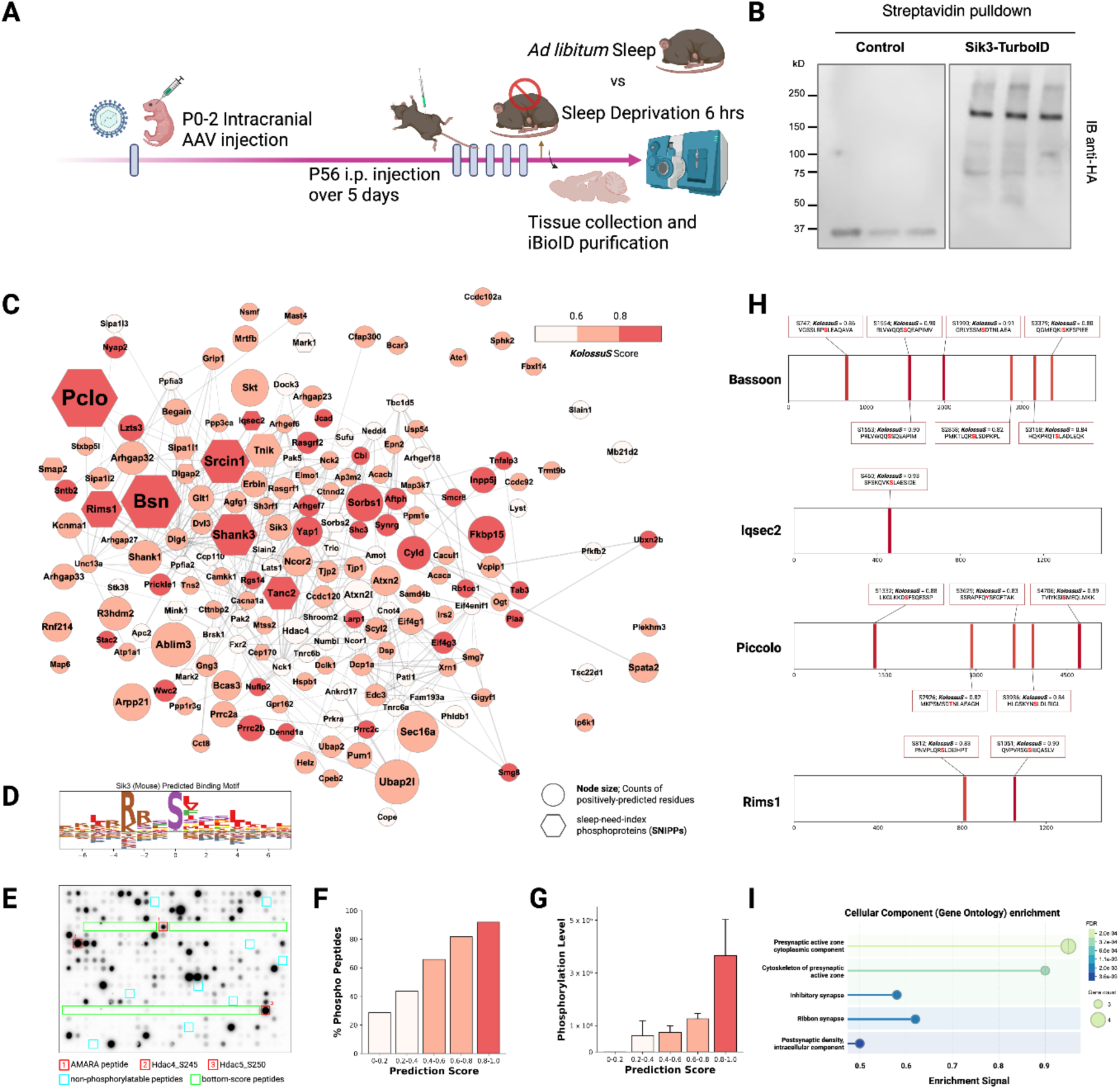
KolossuS and HiUGE-iBioID define a direct substrate network of Sik3 under sleep deprivation. **(A)** Schematic illustration of the HiUGE-iBioID proximity phosphoproteomics workflow used to identify Sik3 substrates in vivo. **(B)** Western blot analysis showing TurboID-HA expression and tagging on Sik3 in mouse brains, compared to a soluble TurboID-HA control expressed via HiUGE-IRES-TurboID. **(C)** Substrate network of Sik3, composed of candidates identified by HiUGE-iBioID and filtered by KolossuS prediction score (≥ 0.50). Node color reflects KolossuS score, and node size indicates the number of unique phosphorylated residues positively predicted by KolossuS. Edges represent STRING-derived protein interactions with medium confidence (0.40). Hexagonal nodes denote sleep-index phosphoproteins (SNIPPs). **(D)** Sequence motif of Sik3 substrates, generated by aligning KolossuS-predicted substrates (KolossuS ≥ 0.75) against the mouse proteome. **(E)** Kinase assay using peptide arrays representing KolossuS- and HiUGE-iBioID–selected candidate substrates. Rectangular highlights indicate: red— positive control peptides, cyan—non-phosphorylatable controls, green—negatively predicted peptides (KolossuS < 0.2). **(F)** Phosphorylation frequency of arrayed peptides, binned by KolossuS score (bin width = 0.2), calculated as the fraction of phosphorylated peptides per bin. **(G)** Phosphorylation intensity of arrayed peptides, averaged within each KolossuS score bin (bin width = 0.2). **(H)** Representative presynaptic Sik3 substrates identified under sleep deprivation. **(I)** Cellular Component Gene Ontology (GO) analysis of high-confidence Sik3 substrates (KolossuS > 0.80), with enriched terms clustered using the Jaccard index via STRING API (https://string-db.org/).

Following proximity labeling, proteins proximal to Sik3 were isolated using streptavidin affinity purification, and phosphorylated peptides were enriched using TiO₂ chromatography as above. LC-MS/MS analysis revealed 2,381 phosphorylated peptides derived from 356 proteins in sleep-deprived samples, compared to only 258 peptides from 168 proteins under ad libitum conditions (Fig. S14, consistent with prior work showing increased phosphorylation during sleep deprivation. Phosphopeptides exhibiting a fold-change ≥1.5 and p-value <0.05 (two-tailed heteroscedastic t-test) compared to the soluble TurboID were considered specific to the Sik3 bait. As we previously observed for other kinases above, 30.14% of the identified phosphopeptides were not listed in the PhosphositePlus database, again highlighting proximity phosphoproteomics ability to provide deep coverage of kinase substrates that may be missed by shotgun analysis of lysates. This experimental approach allowed us to unravel specific in vivo phosphorylation networks associated with Sik3 under conditions of sleep deprivation, providing insights into potential molecular mechanisms underpinning sleep homeostasis.

### HiUGE-iBioID revealed Known and Novel Sik3 Substrates

Previously established Sik3 substrates—including Hdac4, Hdac5, Deptor, Rptor, and Rictor (*88–90*), were specific to Sik3 under both sleep deprivation and ad libitum conditions. This further validates that the HiUGE-iBioID proximity-phosphoproteomics approach can successfully capture known kinase substrates *in vivo*. As before, several other kinases were also detected in proximity to Sik3, such as Camkk1, Pak5, Mark2, and Mast2 (Fig. S14). Many of these also mediate phosphorylation in neuronal synapses (*66, 91, 92*). This complexity again precluded direct attribution of phosphopeptides to Sik3, so we applied the KolossuS model to predict direct Sik3 phosphorylation targets. Of the 1566 detected peptides, KolossuS predicted 523 as likely Sik3 substrates (KolossuS score >0.50, Fig. 5C,D) and 1,010 as unlikely (KolossuS score ≤0.50). KolossuS notably correctly predicted phosphorylation of the established Sik3 target residue Ser245 on Hdac4 (KolossuS = 0.596), while accurately excluding Ser265 (KolossuS = 0.306), which is known to be phosphorylated by PKA (*88, 89, 93*).

### Experimental Validation of KolossuS Sik3 Positive and Negative Predictions

To experimentally validate KolossuS predictions, we synthesized a peptide array consisting of 459 peptides derived from the Sik3 HiUGE-iBioID proximity-phosphoproteomics dataset, including peptides across the KolossuS probability range of Sik3 substrates. In vitro kinase assays using purified Sik3 confirmed phosphorylation of 259 out of 365 positively predicted (KolossuS score >0.5) peptides (70.96%), whereas only 7 of the 46 KolossuS negative peptides (15.22%) were phosphorylated (Fig. 5E). The validation confirmed that KolossuS is well-calibrated, i.e., the distribution of its probability scores matches the observed frequency of phosphorylation events (Fig. 5F). Beyond just the binary measurement of phosphorylation, the level of Sik3-mediated phosphorylation was demonstrably higher for peptides in the 0.8-1.0 probability range (Fig. 5G). Together, these data demonstrate KolossuS correctly learned the peptide substrate features corresponding to the probability of phosphorylation. Furthermore, peptides in the 80-100% probability range are highly phosphorylated and thus likely represent preferred substrates.

### KolossuS-Predicted Sik3 Substrates Define Synaptic Processes in Sleep

Our proximity-phosphoproteomics identified Sik3-proximal proteins phosphorylated during sleep deprivation, including 35 previously described sleep-need-index phosphoproteins (SNIPPs)(*87*) (Fig. 5C). Of these, 18 proteins had residues positively predicted as direct substrates of Sik3 by KolossuS, including Sipa1l1, Bsn, Pclo, Tanc2, and Rims1. Additionally, 14 HiUGE-iBioID-KolossuS substrates were previously identified as Sik3 interactors via co-immunoprecipitation mass spectrometry from earlier studies, such as Amot, Arhgap32, Unc13a, and Git1 (*87*). Gene Ontology (GO) analysis of the 175 proteins predicted as Sik3 substrates (KolossuS >0.5) revealed a significant enrichment in synapse-related terms, particularly postsynaptic density (GO:0014069, FDR = 2.18e-8), represented by SNIPPs proteins including Shank3, Srcin1, Tanc2, Dlgap2, and TNIK. Notably, molecular function terms related to GTPase regulator activity were also significantly enriched (GTPase regulator activity, GO:0030695, FDR = 3.03e-7; GTPase activator activity, GO:0005096, FDR = 0.00090), exemplified by SNIPPs such as Sipa1l1, Trio, and Smap2. Further refinement of GO analysis specifically on the 39 substrates with KolossuS scores greater than 0.8 revealed even stronger enrichment in presynaptic components, presynaptic active zone cytoplasmic components (GO:0098831, FDR = 0.0041), featuring key synaptic SNIPPs proteins such as Rims1, Bsn, Pclo, and Iqsec2 (Fig. 5H,I). These findings collectively reinforce the likely role of Sik3 kinase activity in modulating synaptic processes underlying sleep homeostasis. By integrating KolossuS predictions with altered sleep HiUGE-iBioID phosphoproteomics, we not only identified direct kinase-substrate relationships but also generated a refined map of the local kinase signaling network at higher spatial resolution.

### New Potential Functional Insights into Sik3’s Roles

Proteome-wide computed structural analysis of protein phosphorylation regulatory sites find they are often biased to short disordered regions between folded domains or within domain loops (*94*). Thus, we also examined the sites of Sik3 phosphorylation to identify new potential mechanisms of Sik3 regulation (Table S5). Multiple Sik3 substrates were identified within domains essential for GTPase regulation, including phosphorylation events in the PH2 domains and N-terminal Ras-GEF domains of RasGrf1 and RasGrf2. These sites, conserved across both proteins, suggest an evolutionary pressure towards Sik3 regulation of Ras signaling. Another finding was Sik3-mediated phosphorylation of the SNIPP protein TNIK at residues S649 and S678. These sites in TNIK are within a region known for its interaction with Nedd4 (an E3 ubiquitin ligase). Interestingly, Sik3 also phosphorylated Nedd4 within the TNIK interacting interface at residue S309, suggesting Sik3 might coordinate regulatory interactions within the Nedd4-TNIK signaling axis, perhaps at synapses. NMDA receptor activation can modulate TNIK phosphorylation (*95*), while reducing TNIK levels lowers the surface levels of GluA1 (*96*), suggesting a role in postsynaptic signaling at excitatory synapses. Additionally, TNIK appears to be involved in regulating dendritic outgrowth, likely through its interactions that includes the E3 ubiquitin ligase Nedd4 and Rap2A (*97*). Other intra-domain localized Sik3 substrate sites include the SH2 domains of Shc3 and Nck2, the guanylate kinase-like domain of Tjp1 (two phosphorylation sites), the PDZ domain of Pclo, the PH domain of ArhGef6, and the Mu Homology Domain (MHD) of Ap3m2. These sites suggest Sik3 may broadly broadcast sleep need across a wide network of signaling pathways.

## Discussion

Predicting which proteins a kinase will phosphorylate remains a fundamental challenge in cell signaling research. Traditional approaches to model kinase specificity suffer from several critical limitations. Position-specific scoring matrices (PSSMs) derive scores by evaluating the likelihood of particular amino acids appearing at specific positions around the phosphorylation site, treating each position independently. While this approach has provided a foundation for identifying substrate motifs, its inherent assumption of positional independence fails to capture the complex interdependencies among residues that govern kinase specificity.

Protein language models offer a powerful solution through their rich representation learning capabilities. Trained on millions of protein sequences, these representation models capture evolutionary patterns that encode key structural and functional relationships without requiring explicit structural data. Their self-learned grammar of protein sequences detects subtle interdependencies between amino acid positions, providing an ideal foundation for modeling the complex grammar of kinase-substrate recognition.

KolossuS harnesses these representation learning capabilities by implementing a co-embedding architecture that places kinases and substrates in the same mathematical space. This approach embeds a powerful biological hypothesis—kinases with similar representations will phosphorylate similar substrates—creating a conceptual framework that naturally captures functional relationships across the kinome. Unlike Phosformer-ST, which focuses exclusively on S/T kinases and has uneven performance across kinase families, KolossuS achieves superior accuracy across the entire kinome with well-calibrated probability scores that directly translate to reliable substrate predictions in laboratory settings.

The interpretability of KolossuS embeddings reveals informative patterns of functional organization that we contrasted with kinase evolutionary relationships. We observed a clear separation between the CMGC family of kinases and other S/T kinase families, reflecting their distinctive proline/arginine-directed phosphorylation patterns and unique kinase domains. While KolossuS-derived embeddings broadly agree with phylogenetic groupings, we also observed significant divergence within certain families like AGC and STE, suggesting substantial substrate specificity evolution despite conserved catalytic domains, which has been observed for some STE kinases (*98*). Intriguingly, we also identified convergent evolution across distinct kinase families. The close pairing of DMPK (a Ser/Thr kinase) and LMTK3 (a Tyr kinase) exemplifies this phenomenon, with both recognizing similar cytoskeletal regulatory substrates despite different evolutionary origins. These patterns suggest a spectrum of dual-specificity potential across kinases, a possibility that will require extensive validation as it challenges the traditional dichotomy of tyrosine and serine/threonine kinases.

Position-specific entropy analysis confirms that KolossuS captures crucial positional information despite using average-pooled embeddings. This enables generation of interpretable motif logos for understudied “dark kinases,” providing the first comprehensive atlas of substrate preferences across the entire human kinome. Validating our approach, these logos closely match the established logos of well-characterized kinases like PKA (Ser/Thr kinases), ABL1 and SRC (Tyr kinases). Kinases located near each other in our dendrogram displayed similar motif characteristics, further confirming that mathematical proximity in our embedding space reflects biological reality.

Our design of KolossuS presents some limitations as well as directions for future research. The current focus on 15-mer substrate peptides, while computationally efficient and aligned with available data, may not capture the full protein context that influences in vivo specificity. Future work could also more explicitly incorporate protein structure information through tokenized representations (*99, 100*). Our ablation studies also revealed interesting scaling properties that diverge from reported limitations to PLM scaling. KolossuS continued to improve with larger ESM models (ESM-2 15B vs ESM-2 650M or ESM-C 6B). While the scaling of PLMs has previously been reported to offer diminishing returns when predicting function (*43*), our findings suggest that scaling may indeed be useful in the case of phosphorylation prediction. Moreover, it indicates that our architecture effectively leverages richer representations and could be applied in other settings. In particular, KolossuS’ transfer learning framework can be adapted to any new foundation model and extended to other post-translational modifications with similar sequence-dependent specificity patterns.

Beyond in silico assessments of kinase specificity, kinase-substrate reactions also leverage the critical feature selection of proximity within cells, an attribute that nature has leveraged heavily by use of scaffolding and anchoring proteins, membrane and organelle targeting, docking motifs and interactions, and active transport. These features are poorly understood and currently require experimental approaches. Also, because kinase-substrate interactions are transient and are influenced by endogenous protein levels, this has been a challenging domain to address.

To address these significant barriers, we incorporated proximity proteomics with Kolossus predictions. Specifically, we leveraged a recent gene editing approach we developed to introduce TurboID into endogenous kinases in vivo, which covalently marks nearby proteins. We reasoned this would effectively enable the capture of even transient interactions. No approach is perfect and one unavoidable limitation to this, and all fusion-protein approaches, is that bait functions may be altered by the fusion. We have previously demonstrated that careful choice of the fusion sites can insert TurboID into sites that minimize the deviation of the computed folded structure from native state, and have validated that our designs can recapitulate known interactions (*40*) and kinase substrates (described above).

A challenging aspect of substrate profiling is that phosphorylation is cell-state dependent and transient and thus reported kinase substrates can differ between studies. Thus, analysis of in vivo reactions by HiUGE-iBioID-kinase can highlight diverse substrates across distinct conditions. For example, by assaying under both normal and sleep-deprived conditions, we could study Sik3, a kinase pivotal to sleep homeostasis and which is known to be activated by sleep deprivation.

With sleep deprivation, HiUGE-iBioID and KolossuS identified several proteins were heavily phosphorylated by Sik3 in vivo that may explain mechanisms of how it modulates sleep. One interesting finding was the conserved Sik3 phosphorylation sites shared between RasGrf1 and RasGrf2 GEF and PH domains. Prior studies have shown that RasGrf1 is upregulated by light within the suprachiasmatic nucleus (SCN) and that Ras activity also shows circadian oscillations in the SCN. Importantly, Sik3 has now been shown to act within SCN neurons, and conditional loss of Sik3 in the SCN perturbs circadian locomotor rhythms (*101*). These findings, together with the role of Ras signaling in the SCN, suggest that Sik3-dependent phosphorylation of Ras pathway components may fine-tune sleep regulation. In a recent study, Mikhail et.al. (*102*) showed ERK phosphorylation levels, which is downstream of Ras, vary with sleep and wakefulness, with increases during wakeful periods and decreases during sleep. Deletion of the Erk1 or Erk2 genes led to a notable increase in the duration of wakefulness, while small molecule inhibition of ERK phosphorylation led to a reduction in sleep time and extended the length of wakefulness bouts.

The most highly phosphorylated proteins were the large presynaptic proteins piccolo and bassoon, which together govern the assembly and maintenance of the presynaptic active zone, and may modify the presynaptic cytoarchitecture through regulating activity-coupled ubiquitination or autophagy (*103, 104*). The large number of Sik3 phosphorylation sites on both proteins (6 sites on Bassoon, 5 on Piccolo) highlights the potential for Sik3 to dramatically modulate their function. Modulation of the active zone in relation to sleep pressure appears to be evolutionarily conserved. For example, regulation of the levels of Bruchpilot (BRP) in drosophila, a large presynaptic scaffolding protein, also regulates sleep need (*105*). Recent evidence suggests this process may also link aging and sleep disturbances (*106*).

Transitions between wakefulness and sleep involve rapid reconfigurations of cortical network activity. Emerging research indicates that the ability to switch states also relies on proper presynaptic neurotransmission. A striking example is the discovery of the “restless” mouse mutant (rlss), which carries a mutation in the vesicle-release protein synaptobrevin-2 (VAMP2). This mutation causes a global reduction in presynaptic neurotransmitter release probability, effectively dampening synaptic strength across the brain (*107, 108*). Intriguingly, rlss mice show severe abnormalities in state transitions: they stay awake for abnormally long bouts and then crash into long bouts of sleep, with far fewer brief transitions between states. Finally, several Sik3 substrates in the sleep deprived state were also found modulate postsynaptic functions (Shank3, Srcin1, Ablim3, Tanc2, TNIK, and NEDD4). Recent work, using modeling as well as chemogenetics to induce postsynaptic potentiation, demonstrates that elevated synaptic strength of prefrontal cortical excitatory neurons drives sleep need (*109*). Our combined experimental and computational results suggest Sik3 may serve as central signaling node to orchestrate both pre and postsynaptic activity through the substrates identified.

## Conclusion

In conclusion, kinase signaling has been extensively studied, yet the combinatorial and complex factors of proximity in cell space and phosphoacceptor recognition preferences have stymied efforts to identify substrates in physiological settings. Our combined protein language-based KolossuS model and HiUGE-BioID-kinase analysis addresses these bottlenecks and allows for analysis of data from in vivo settings, even under different physiological states. When applied to sleep, it revealed both known and new in vivo substrates of Sik3 in relation to sleep need. The generalizability of HiUGE-iBioID across proteins (40, 71) and the broadly learned kinase-substrate features of KolossuS present a combined approach that promises to reveal new insights into kinome function. Our results point to multiple new directions to deconvolve other post-translational signaling pathways as larger datasets for training become available. We also anticipate KolossuS will continue to improve by incorporating either future PLM foundation models and/or by learning based on full-length substrates to infer co-localization mechanisms based on sequence driving subcellular localization (refs for PLM prediction of localization). Finally, our results call for a deeper integration of machine learning models and experimental techniques to enable discoveries that would be out of the reach of either approach individually.

## Declarations

This work was supported by National Institute of Mental Health Grant R01MH126954 and in part by grant 2023-332396 from the Chan Zuckerberg Initiative DAF, an advised fund of Silicon Valley Community Foundation to S.H.S.; and the Whitehead Scholarship to R.S. We thank Dr. Erik Soderblom (Duke Proteomics and Metabolomics Shared Resource), Sean Gay and Julie Kent for technical support and advice. D.S. is supported by a PhRMA Foundation Predoctoral Fellowship in Drug Discovery. A.P. is a Duke AI Health Data Science (AI-HDS) Fellow. AI-HDS at Duke is supported by the National Center for Advancing Translational Sciences (NCATS), National Institutes of Health, through Grant Award Number UL1 TR002553. The Duke AI Health Data Science Fellowship Program is supported by the above grant, the Duke Department of Biostatistics & Bioinformatics, and Duke AI Health. The content of this publication is solely the responsibility of the authors and does not necessarily represent the official views of the NIH. This work was also supported by Japan Society for the Promotion of Science KAKENHI Grant-in-Aid for Scientific Research (22H04918 and 22K21351 to M.Y., 23KJ0269 to H.Y.), Japan Science and Technology Agency Strategic International Collaborative Research (JPMJSC2204 to T.F.), Japan Agency for Medical Research and Development under Grant Number JP21zf0127005 to M.Y., and World Premier International Research Center Initiative from Ministry of Education, Culture, Sports, Science and Technology to M.Y.

