## Supplemental methods and data for "Deep Learning-coupled Proximity Proteomics to Deconvolve Kinase Signaling In Vivo"

### Materials and Methods

#### Data curation and pre-processing

We integrated three complementary datasets of kinase-substrate interactions, each offering different trade-offs between coverage and in vivo fidelity. The broadest coverage comes from “Atlas” datasets that used positional scanning peptide arrays (PSPAs) to estimate position-specific scoring matrices (PSSMs) for 303 Ser/Thr kinases and 92 Tyr kinases. These PSSMs were then screened against sub-sequences of the human proteome to estimate kinase-specific binding strength of 11-mer peptide sequences. While comprehensive, these data represent in vitro estimations. The next tier of data comprises Sugiyama et al.'s "Human Kinome" dataset, where dephosphorylated cell lysates were assayed against kinases, offering greater biological context. The highest fidelity but smallest dataset comes from literature-curated interactions in the Phosphosite database, documenting specific substrate proteins and their phosphorylation sites validated in vivo or in controlled in vitro studies. Kinases that could not be assigned to a hand-curated kinase-family mapping（sourced from (*1*) and Uniprot）were removed.

To construct positive and negative labeled examples from the Atlas datasets, we designated all substrate 15-mers with a Ser/Thr kinase PSSM percentile score >= 95 (or <= 5) as a positive (or negative) example for that kinase. For Tyrosine kinases, the positive (negative) cutoff scores were >= 90 (<= 10). For the Human Kinome and Phosphosite datasets, positive interactions were already provided; negatives were created by pairing the kinase with a substrate that was not observed to be phosphorylated by that kinase or any of its homologues to create datasets with a 50:50 ratio of positive to negative examples. We note that the issue of curating appropriate negatives for the latter two datasets is a difficult task, due to the lack of existing data on true negatives for kinase-substrate interactions. Several options were considered, including the use of random sequences for negatives. However, due to our desire to train the model on “hard negatives”-- i.e., sequences that are not phosphorylated by the kinase under consideration, but are known to be phosphorylatable-- we decided to take the approach described above.

To ensure robust evaluation of generalization, we constructed consistent train-validation-test splits across both the atlas and human kinome datasets. Rather than using sequence similarity thresholds, we split the data based on kinase families. This choice reflects the deep evolutionary relationships captured by kinase family annotations, which incorporate not just sequence similarity but also evolutionary similarity, based on phylogenetic analyses. Specifically, we held out 2 kinase families (including CK1, JAK kinases) for validation and 2 families (TKL, FAK kinases) for testing. Additionally, within each family in the training set, we held out 16 individual kinases for validation and 16 for testing. This two-level holdout strategy allows us to evaluate generalization both within and across families. The same families and individual kinases were held out across both datasets, ensuring that our evaluation accurately captures the model's ability to generalize across evolutionary distances. As a third tier of evaluation, 7.5% of randomly sampled observations were held out for validation and test independently to assess kinase affinity across substrates. The Phosphosite database was reserved entirely for final evaluation, providing an independent assessment on high-confidence in vivo interactions.

**Model Architecture**

The KolossuS architecture is molded to these data characteristics. The model accepts a full kinase sequence but processes substrates as 15-mer subsequences-- the minimum context shared across our datasets. This design choice can leverage the comprehensive PSPA data while maintaining compatibility with the other datasets. We employ the ESM-2 protein language model to generate initial embeddings, capturing both local substrate sequence patterns and global kinase protein context. In ablations, we evaluated both the commonly-used 650-million parameter model (with 33 transformer layers) and the largest 15-billion parameter model (with 48 transformer layers). The latter consistently outperformed the former.

The architecture comprises two key components. The sequence embedding module processes kinase and substrate sequences separately through parallel feed-forward networks, each transforming the 5120-dimensional ESM-2 embeddings to a shared 1024-dimensional space. This projection preserves sequence-structure relationships while enabling direct comparison between kinases and substrates. The prediction module then computes the probability of phosphorylation using cosine similarity between the projected kinase and substrate embeddings. This co-embedding approach not only provides accurate predictions but also enables direct analysis of kinase-substrate relationships in the shared latent space.

**Model Training**

We employed a two-phase training approach to leverage the varying quality of our datasets. Initial training used the broad PSPA data to learn general sequence-structure relationships. This was followed by fine-tuning on the human kinome dataset to capture more biologically-relevant patterns. The Phosphosite database was held out entirely for evaluation.

For model assessment, we computed both threshold-independent metrics (area under ROC and precision-recall curves) and fixed-threshold metrics (accuracy, sensitivity, and specificity at probability threshold 0.5). While AUC metrics are standard for machine learning evaluation, biological applications require predictions to be well-calibrated across diverse kinase families. We therefore emphasized performance at a fixed threshold, evaluating consistency across different kinase families and evolutionary distances.

**Implementation Details**

The model was implemented in PyTorch and trained on NVIDIA A100 GPUs. ESM-2 embeddings were extracted from the final layer, excluding padding and start-of-sequence tokens. The feed-forward networks use ReLU activation and Xavier-initialized weights. We trained using the AdamW optimizer with learning rate 1e-4 and binary cross-entropy loss. The initial training phase ran for 60 epochs, followed by 20 epochs of fine-tuning. Model selection was based on validation performance.

**Computational Requirements**

The model contains approximately 25 million trainable parameters. Training completes in approximately 5.5 hours on a single A100 GPU, with inference taking ~0.0004 seconds per kinase-substrate pair. This efficiency enables rapid screening of potential substrates across the proteome.

#### Animal

All of the *in vivo* experiments were conducted at Duke University. For in vivo CRISPR-Cas9-related experiments, H11-Cas9 mice (Jackson Laboratory #28239) were used. Mice were housed under proper conditions in the Division of Laboratory Animal Resources facility at Duke University, following the approved guidelines of the Duke University Institutional Animal Care and Use Committee (Protocol #A144-23-07). Mice were maintained under 72 ± 2 °F ambient temperature, 30-70% humidity, and a 12-hour light/dark cycle.

#### AAV Preparation

HiUGE TurboID knock-in plasmids previously described ((*2*), Addgene Plasmid #200383, #200384, #200385) were used to endogenously tag six kinases—Sik3, Aak1, Akt1, Brsk2, Camk1d, and Uhmk1—with HA-tagged TurboID. Briefly, a donor of HA-tagged TurboID coding sequence was flanked by DNA sequences specifically recognized by a synthetic donor-specific gRNA (DS-gRNA), which is inert to the C-terminal of each target gene. For each kinase, a gene-specific gRNA (GS-gRNA) expression cassette was inserted in tandem with the DS-gRNA expression cassette, allowing single-vector delivery of both components. GS-gRNAs were designed using CRISPOR, and a pair of 23–24 nucleotide oligonucleotides were annealed and ligated into the SapI site of the GS-gRNA cassette. The gRNA sequences used were as follows: Sik3; TTCCATATCTTGCTACACGC, Aak1; CCTAGCCAGGTCTTTGCTGC, Akt1; CAGTTCTCCTACTCAGCCAG, Brsk2; CACCGAGTACCCGATGGGCA, Camk1d; AGGCCCACCACTGTGACAAC, and Uhmk1; CCGCTGAGTGCCTACAAGAG. To endogenously express soluble TurboID as a background detection control, C-terminal regions of the target genes were modified using a donor containing a stop codon followed by an internal ribosome entry site (IRES) and the TurboID-HA coding sequence.

AAV was prepared following previously described methods (*2*). Briefly, concentrated AAV virus was produced in HEK293T cells grown in six 15-cm dishes by triple transfection with 15 μg HiUGE vector, 30 μg pADdeltaF6, and 15 μg serotype plasmid (pUCmini-iCAP-PHP.eB, a gift from Viviana Gradinaru, Addgene plasmid #103005). Three days post-transfection, the cells were lysed and the virus was concentrated using an Optiprep density gradient (Sigma #D1556). For small-scale AAV production, HEK293T cells were grown in 12-well plates and transfected using 0.4 μg HiUGE vector, 0.8 μg pADdeltaF6, and 0.4 μg serotype 2/1 plasmids. Three days post-transfection, the virus-containing medium was filtered through Costar Spin-X columns (Sigma #8162) and stored at 4°C until use.

#### AAV Injection and Sleep Deprivation for Proximity-phosphoproteomics

Neonatal (P0-2) H11-Cas9 mice were anesthetized by hypothermia and injected intracranially with purified AAV (2 μL per hemisphere, PHP.eB serotype, >10^10 GC/μL titer). The negative control used was an IRES-TurboID donor HiUGE-construct. At 54-58 days post-AAV injection, mice received daily intraperitoneal (i.p.) injections of 5 mM biotin (50 mg/kg, 0.5 mL) for 5 consecutive days. One day following the fifth injection (P59-63), mice received biotin injections at the onset of the light phase (ZT0) and were subsequently subjected to sleep deprivation for

6 hours through cage changing, intermittent mechanical agitation, and the introduction of novel enrichment items. Food and water were available throughout the sleep deprivation period (*3*). Immediately following sleep deprivation, cortices and hippocampi were harvested, flash-frozen in liquid nitrogen, and stored at -80°C for downstream processing.

**Proximity-phosphoproteomics Sample Purification**

For each replicate, cortex and hippocampus tissue from three mice was collected, flash-frozen, and combined. The tissue was homogenized using a Dounce homogenizer in RIPA buffer (150 mM NaCl, 50 mM Tris-HCl, 0.1% SDS, 1% Triton X-100, 1% sodium deoxycholate, 1 mM EDTA) supplemented with cOmplete protease inhibitor cocktail (Roche #11836145001) and PhosSTOP phosphatase inhibitor cocktail (Roche #04906845001). Homogenates were sonicated for three 10-second intervals and centrifuged at 14,000 × g for 30 minutes at 4°C. The resulting supernatant was desalted using Zebra Spin Desalting columns (7K MWCO, Thermo Fisher #89892). The flow-through was combined with 50 µL Pierce Streptavidin Magnetic beads (Thermo Fisher #88817) and rotated overnight at 4°C.

The following day, beads were transferred to low-protein-binding tubes and washed twice with RIPA buffer, once with 1 M KCl, once with 0.1 M Na2CO3, once with 2 M urea in 10 mM Tris-HCl, and twice with RIPA buffer. Biotinylated proteins were eluted by heating the beads in 90 µL of 2× elution buffer (4% SDS, 50 mM Tris, 100 mM NaCl, 20 mM DTT) supplemented with 5 mM biotin. The eluted samples were used for downstream LC-MS/MS and Western blot analysis. For CaMK1D-TurboID samples, beads were additionally incubated in reaction buffer containing 1 mM $CaCl_2$ and 20 ng/µL calmodulin (Sigma #208694-1MG) at 37°C for 30 minutes prior to elution.

**Phosphopeptide Enrichment**

Six tissue samples (3 per condition) were received and stored at −80 °C until processing. Samples were spiked with either 1 or 2 pmol of bovine casein as an internal quality control standard. Samples were then reduced at 80 °C for 15 min, alkylated with 20 mM iodoacetamide for 20 min at room temperature (RT), and supplemented with a final concentration of 1.2% phosphoric acid. After acidification, 559 µL of S-Trap (Protifi) binding buffer (90% MeOH/100 mM TEAB) was added. Proteins were trapped on S-Trap micro cartridges and digested using 20 ng/µL sequencing grade trypsin (Promega) for 1 hr at 47 °C. Peptides were then eluted sequentially with 50 mM TEAB, followed by 0.2% formic acid (FA), and finally with 50% acetonitrile (ACN)/0.2% FA. All samples were subsequently lyophilized to dryness.

For phosphopeptide enrichment, dried peptides were resuspended in 80% ACN/1% TFA and enriched using titanium dioxide ($TiO_2$) tips (GL Biosciences) according to the manufacturer's recommended protocols, employing a competitive modifier to enhance phosphopeptide binding. Enriched phosphopeptides were again frozen and lyophilized to dryness.

**Quantitative LC-MS/MS Analysis**

For LC-MS/MS analysis, enriched phosphopeptide samples were resuspended and loaded onto EvoTips (EvoSep) for automated LC separation using an EvoSep One UPLC system coupled to a Thermo Orbitrap Astral high-resolution accurate mass tandem mass spectrometer (Thermo). Each sample was separated on a 1.5 µm EvoSep C18 column (150 µm × 150 mm) at 55 °C using the SPD30 gradient. Quantitative MS data were acquired using a data-independent acquisition (DIA) method. Full MS scans were performed in the Orbitrap at a resolution (r) = 240,000 (@ m/z 200) from m/z 380–1080 with a target AGC value of 4e5. DIA MS/MS scans were acquired across fixed 5 m/z windows covering m/z 380–1080. MS/MS scans were collected in the Astral analyzer using an AGC target of 5e4, a maximum fill time of 8 ms, and a normalized HCD collision energy of 27%. Each sample injection was completed in approximately 40 min.

Following LC-MS/MS analysis, nine total datasets were imported into Spectronaut (Biognosys), where retention times and precursor/fragment ion masses were aligned across all runs. Relative phosphopeptide abundance was determined using MS2 fragment ion intensities extracted from aligned chromatographic features. Database searching was performed against the SwissProt M. musculus database, supplemented with a common contaminant/spiked protein database (e.g., bovine albumin, bovine casein, yeast ADH) and reversed-sequence decoys for false discovery rate (FDR) calculation. A Spectronaut library was generated exclusively from data acquired within this study. Database search parameters included fixed carbamidomethylation of cysteine, variable oxidation of methionine, variable N-terminal acetylation, and variable phosphorylation on serine, threonine, and tyrosine residues. Trypsin was used as the digestion enzyme, and precursor/product ion tolerances were set to 10 ppm and 20 ppm, respectively. Peptide and protein identifications were annotated at a maximum of 1% FDR based on q-value calculations.

As in the prior method, peptide homology was addressed using razor rules to assign peptides uniquely to the protein group for which they provided the most evidence. Protein homology was addressed by grouping proteins sharing identical sets of peptides and designating a master protein based on percent coverage. Raw phosphopeptide intensity values were exported from Spectronaut for downstream processing. To address missing data points (often due to signal intensities falling below the limit of detection), filters and imputation strategies were applied as previously described. Specifically, phosphopeptides were first required to be detected at least twice across all samples and found in at least 50% of one biological group. Those not meeting this criterion were removed.

For missing values, the following imputation rules were applied: (1) if fewer than 50% of signals were missing within a group, a randomized value near the average of the remaining detected intensities was used; (2) if more than 50% of signals were missing within a group, a randomized intensity within the bottom 1% of detectable signals was imputed. The resulting imputed phosphopeptide intensities were then filtered to include only phosphopeptides with ≥75% localization confidence at the phosphosite(s). Normalization was performed using a robust mean approach. The top and bottom 10% of phosphopeptide signals were excluded, and the mean of the remaining phosphopeptides was used as a normalization factor across all samples. Phosphopeptides passing all criteria and transformations were subjected to statistical analyses, including calculation of fold changes and two-tailed heteroscedastic t-tests on $log_2$-transformed data to identify significant differences between sample groups.

After database searching at a 1% peptide-level FDR, a total of 14,154 phosphopeptides corresponding to 2,173 phosphoproteins were identified. Measures of technical reproducibility, assessed by calculating average %CV across groups, showed that %CVs ranged from ~19.5% for the spiked quality control pool to ~30.5% for other samples, indicative of stable phosphoproteomic enrichment and analysis.

### Phosphorylation of Peptide Array

The peptide array was SPOT synthesized on a cellulose membrane (PepSpot, JPT Peptide Technologies) containing 430 positive peptides (prediction score > 0.5), 47 peptides with low prediction scores (prediction score 0.002-0.034), 9 non-phosphorylatable peptides (derived from positive peptides with phospho-acceptor residues substituted with alanine), and two known Sik3 substrates: HDAC5tide [PLRKTASEPNLKRRR]1 and AMARA peptide [AMARAASAAALARRR].

The membrane was first incubated in methanol for 10 minutes and subsequently rinsed twice with Sik3 kinase reaction buffer (25 mM Tris-HCl, pH 7.5, 0.5 mM EGTA, 0.01% Triton X-100, 5 mM $MgCl_2$, 2.5 mM DTT)4 supplemented with 1 mg/mL BSA. The membrane was then blocked overnight at room temperature in the same buffer. After blocking, the buffer was replaced with fresh blocking solution containing 100 µM ATP, and the membrane was equilibrated at 30°C for 30 minutes.

The phosphorylation reaction was initiated by adding 10 mL of reaction buffer supplemented with 5 µg/mL recombinant Sik3 (S12-11G, SinoBiological) and 50 µCi [$\gamma$-$^{32}$P] ATP (Revvity), and incubating at 30°C for 2 hours with gentle rocking. The buffer was then discarded, and the membrane was washed 10 times with 1 M NaCl, followed by two washes with deionized water. The membrane was subsequently washed with 4 M guanidine hydrochloride, rinsed with deionized water, and washed three times with buffer (50 mM Tris-HCl, pH 7.5, 1% SDS, 0.5% β-mercaptoethanol). Finally, it was rinsed several times with deionized water.

The membrane was placed in a plastic bag, and the incorporation of $^{32}$P into the peptides was visualized by FLA-7000 image analyzer (Fujifilm, Japan). The phosphorylation intensities at each position were quantified using ImageJ. The average intensity of the 9 non-phosphorylatable peptide spots was subtracted for normalization.

### Genome-Wide Kinase-Substrate Prediction

To create a predicted human and mouse phosphoproteome, a set of 466 known human kinases and 457 known mouse kinases were sourced from pkinfam (predicted inactive kinases excluded), a curated list of all human and mouse kinases by Swiss-Prot (https://ftp.uniprot.org/pub/databases/uniprot/current_release/knowledgebase/complete/docs/pkinfam.txt). Substrate sequences were sourced from literature-curated 15-mer peptides that have been experimentally verified to be phosphorylatable (https://www.phosphosite.org). In total, 227,736 and 100,981 substrates were used in the analysis for human and mouse, respectively. KolossuS was then run on the cartesian product of all substrate and kinase sequences. For the kinases, the canonical Uniprot sequences were used.

#### Logo Creation

Putative motifs were predicted for each kinase from the predicted phosphoproteome using the following methods. For each kinase, a weighted position-specific amino-acid count matrix was created from substrates with which the kinase had a positive prediction for phosphorylation (binding score >= 0.75). Counts were weighted by their KolossuS binding scores. The count matrices were then normalized to obtain position-specific amino-acid probabilities. To quantify the degree of residue conservation at each of the non-central (i.e., non-phosphorylated) positions, the relative entropy was computed against the background distribution of position-specific amino acid frequencies computed from the set of all substrate sequences (227,736 human, 100,981 mouse). For the central position, the relative frequency was used, scaled to the maximum relative entropy observed in the non-central position. Logos were visualized using the Logomaker package (version 0.8.6) in Python (version 3.9.0).

#### Kinase Dendrogram Creation

Dendrograms were created from the predicted human phosphoproteome by extracting KolossuS embeddings for each of the kinases in seven selected families, and creating a pairwise similarity matrix using the cosine similarity metric. Similarities were converted to distances by applying the inverse cosine function. The dendrogram was created from the resulting distance matrix using the neighbor-joining method implemented in the R package APE (version 5.8). The dendrogram was plotted using the “treeplot” function of ggplot2 (version 3.5.1). Commands were run on R version 4.2.1.

### Position-specific entropy analysis

For each dataset (Atlas, Human Kinome, and Phosphosite), kinase-substrate pairs that had positive phosphorylation events were extracted. For each kinase, a position-specific amino-acid count matrix was created from its corresponding positive substrates, and counts were normalized to obtain position-specific amino-acid probabilities. The entropy at each position *j* ($j = -7, -6, \ldots, +6, +7$) was calculated using the formula

$$E_j^{(k)} = \sum_{i=1}^{20} p_i * \log p_i,$$

where $E_j^{(k)}$ is the entropy at position *j* for kinase *k*, and $p_i$ is the probability the *i*-th amino-acid. Entropy values were normalized across positions and kinases by subtracting the overall mean and dividing by the overall standard deviation. The distribution across kinases of normalized position-specific entropies was visualized via boxplots (Fig. S8).

### Evaluation of effects of position-specific substrate mutations on KolossuS predictions

For each dataset (Atlas, Human Kinome, and Phosphosite), substrates were “mutated” at the -4 through +4 positions (relative to the phosphorylation site at position 0) by replacing the amino acid at that position by other randomly chosen amino acids. Embeddings were created for this “mutated” substrate for both the ESM-2 15B and KolossuS models, and the cosine distance between this mutant embedding and the original embedding was computed. Cosine distance values were normalized similarly to the normalization procedure used in the position-specific entropy analysis and visualized via boxplots (Fig. S8).

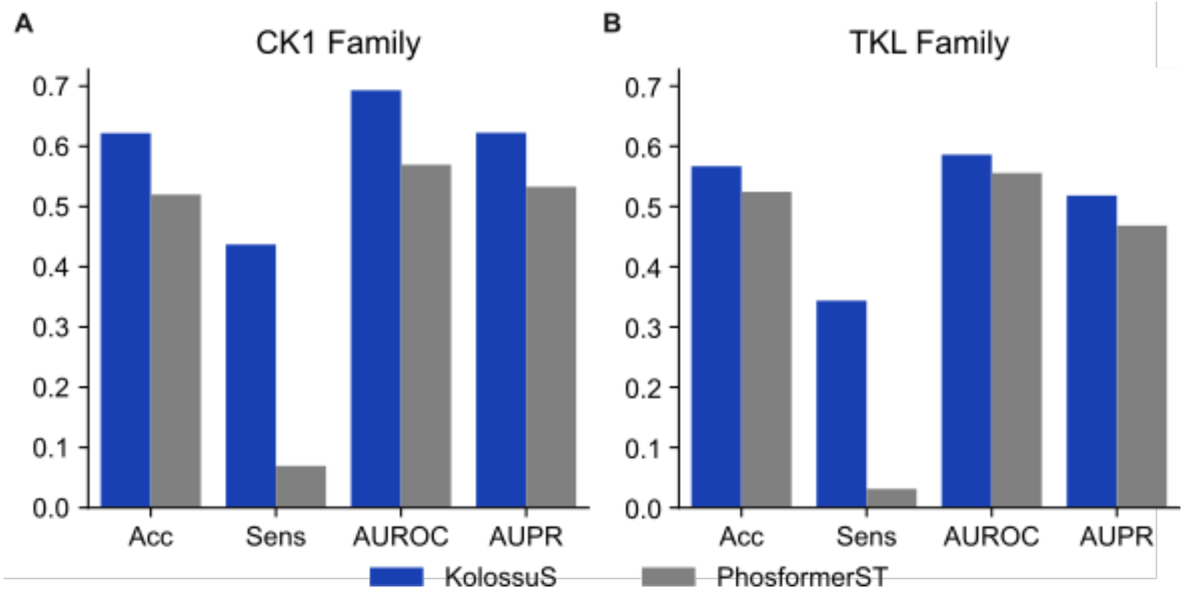


**Fig. S1.** Performance comparison between KolossuS and Phosformer-ST for heldout Ser/Thr kinase families. **(A)** Comparison for the CK1 family. **(B)** Comparison for the TKL family. We note that we used the pre-trained Phosformer-ST model which may not have held out these evaluated kinases from its training data.

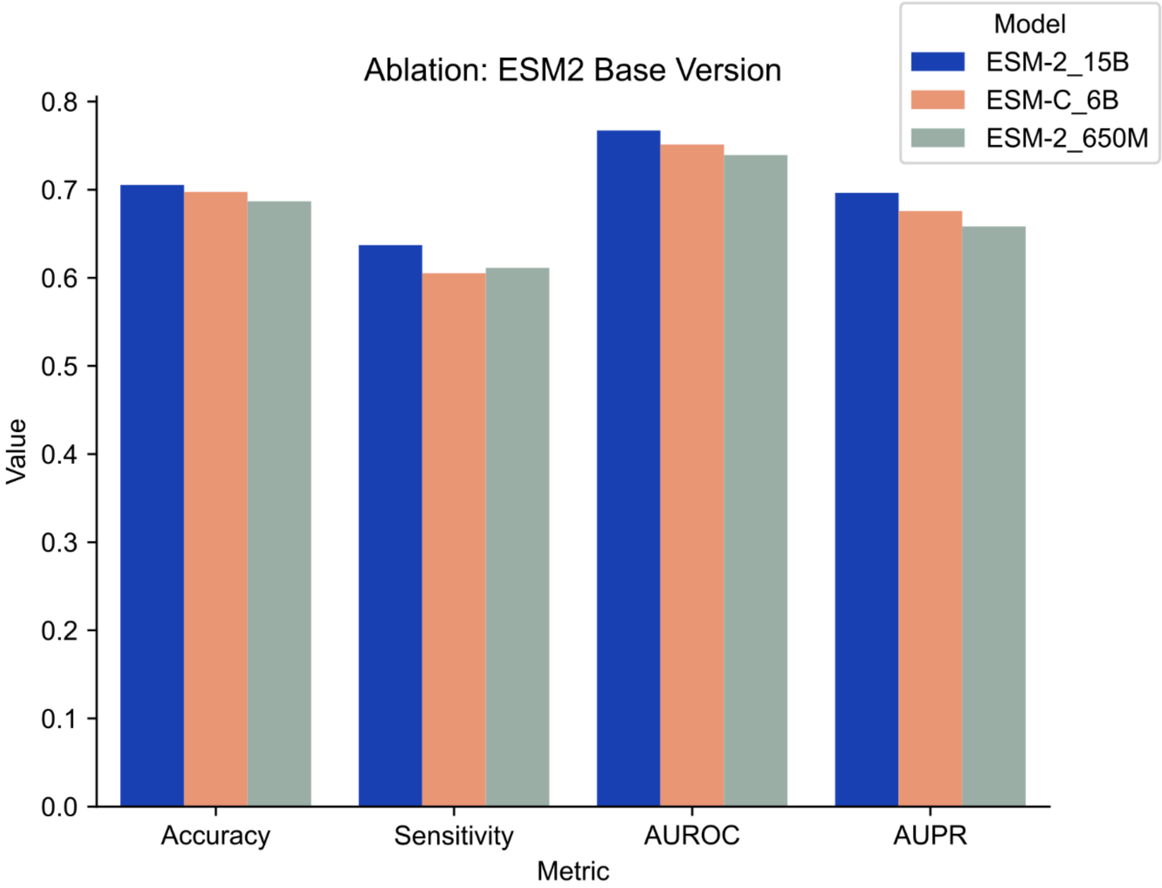


**Fig. S2.** Comparison of classification performance depending on the base version of ESM model (ESM-2 15B, ESM-2 650M, or ESM-C 6B) used to generate embeddings as input to KolossuS. The models were benchmarked on the Phosphosite evaluation dataset, aggregated across human, mouse and rat Ser/Thr and Tyr kinases.

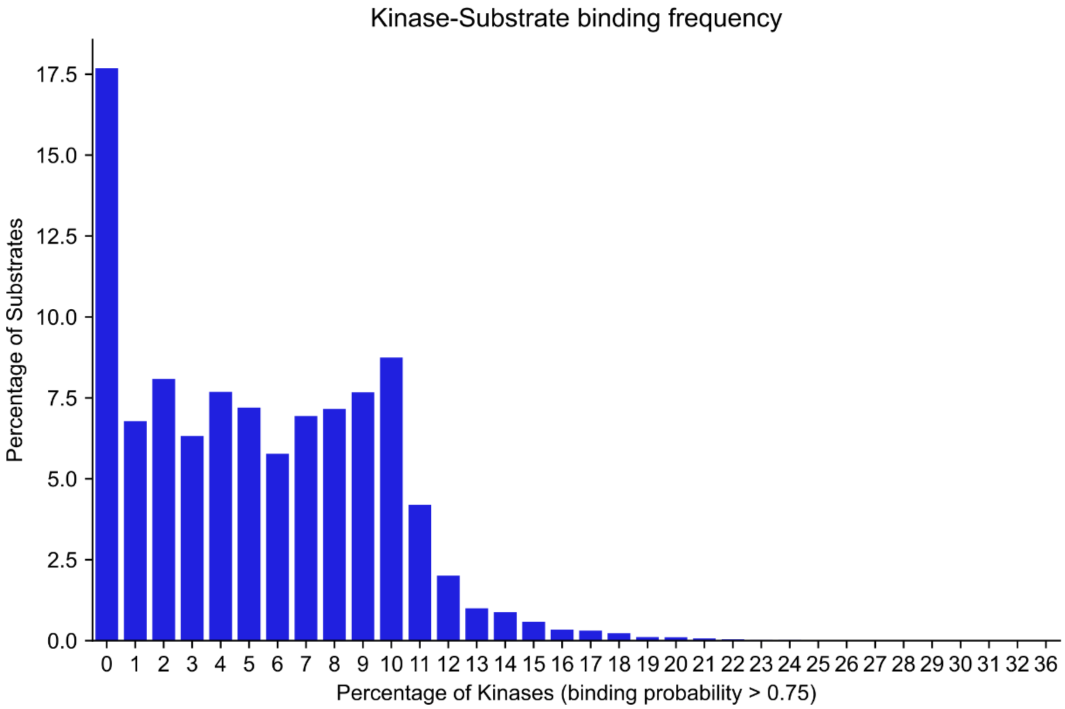


**Fig. S3.** Barplot showing the breakdown of predicted kinase-substrate binding frequencies. The value on the y-axis shows the percentage of substrates predicted to be phosphorylated by *x*-percentage of kinases. For example, 17.5% of substrates are phosphorylated by between 0-1% of kinases, and around 7% of substrates are phosphorylated by 1-2% of kinases. Cumulatively, therefore, over 90% of substrates are predicted to be bound by fewer than 11% of kinases.

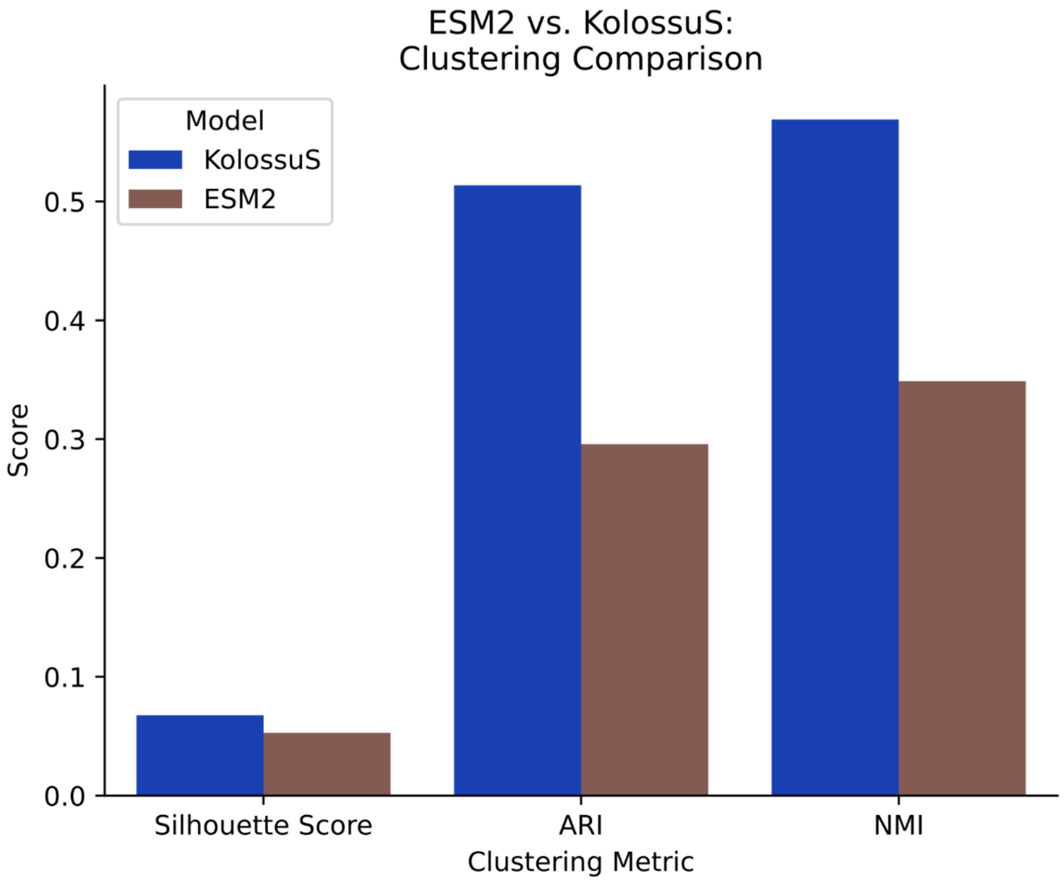


**Fig S4.** Clustering comparison beween KolossuS kinase embeddings and ESM-2 15B kinase embeddings. Three metrics are shown: 1) the silhouette score of the family-label clustering of KolossuS embeddings; and 2) the adjusted rand index (ARI), 3) the Normalized Mutual Information (NMI) between the family-label clustering and a k-means clustering of the embeddings (number of clusters set equal to the number of families). The silhouette score of a clustering of N points is an internal metric of clustering quality, and is calculated by the formula $S = \frac{1}{N}\sum_{i=1}^{N} 1 - \frac{a_i}{b_i}$, where $a_i$is the average distance of the *i*-th to the points within its assigned cluster, and $b_i$is the minimum average distance of the *i*-th point to the points in a cluster that is *not* its own assigned cluster. The Silhouette score can range between $-\infty$and 1; higher scores correspond to well-defined and better-separated clusters. The ARI and NMI are two measures of the average level of agreement between two different clusterings of a set of *N* points; here, the comparison of the family-based clustering to the k-means clustering of the embeddings is an external measure of the extent to how likely two kinases whose KolossuS-based representations have a small distance from each other are to be from the same family, and vice versa. The centroids of the k-means clusterings were initialized as the centers of mass of the clusters defined by the family labels. Taken together, these results suggest that the KolossuS embeddings better agree with the phylogenetic grouping of kinases than ESM-2 embeddings.

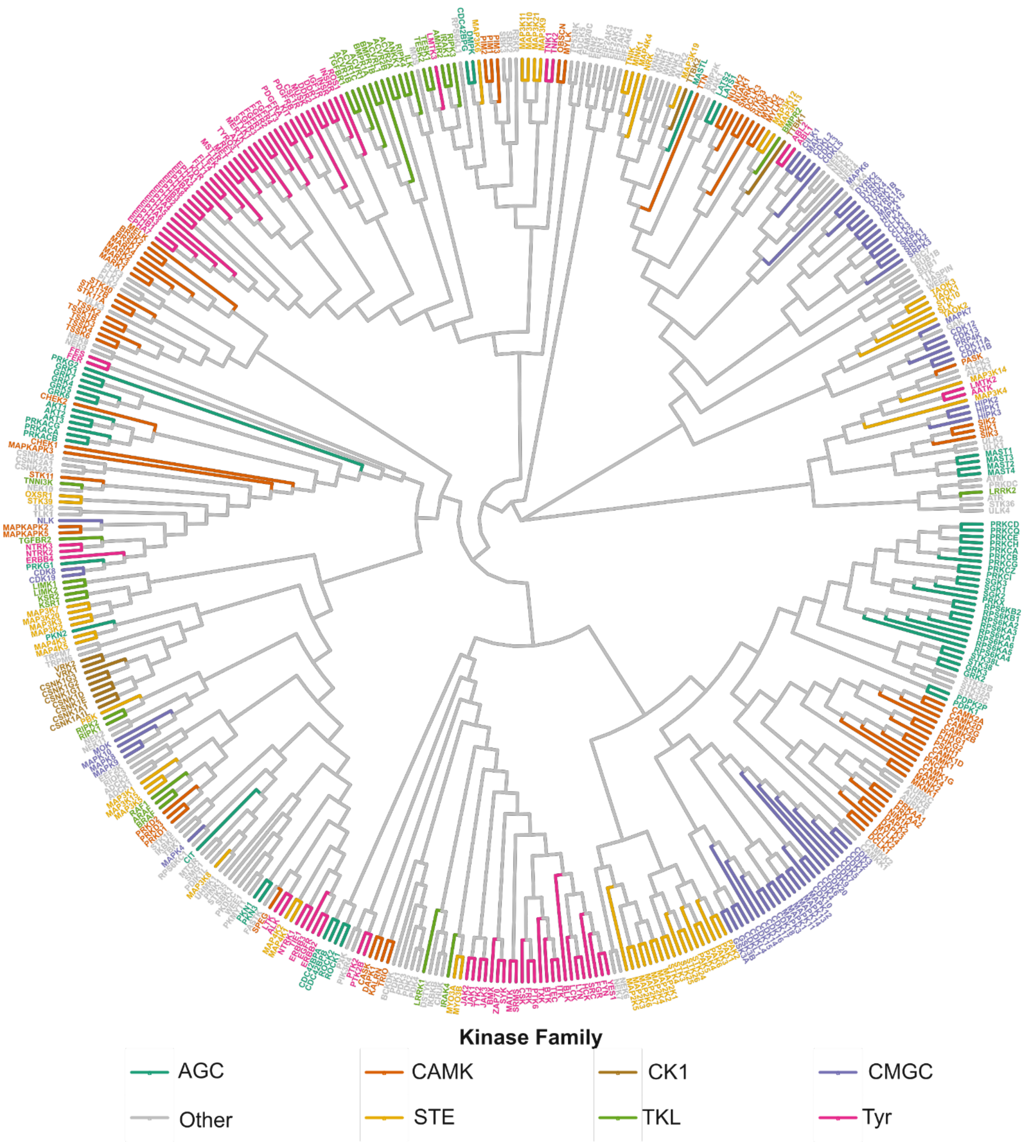


**Fig. S5.** Dendrogram of the 466 kinases used in the genome-wide kinase-substrate phosphorylation prediction, based on raw ESM2 15B protein language model embeddings. The same methodology was used to generate the dendrogram as the KolossuS embedding-based Dendrogram in Fig. 4(D).

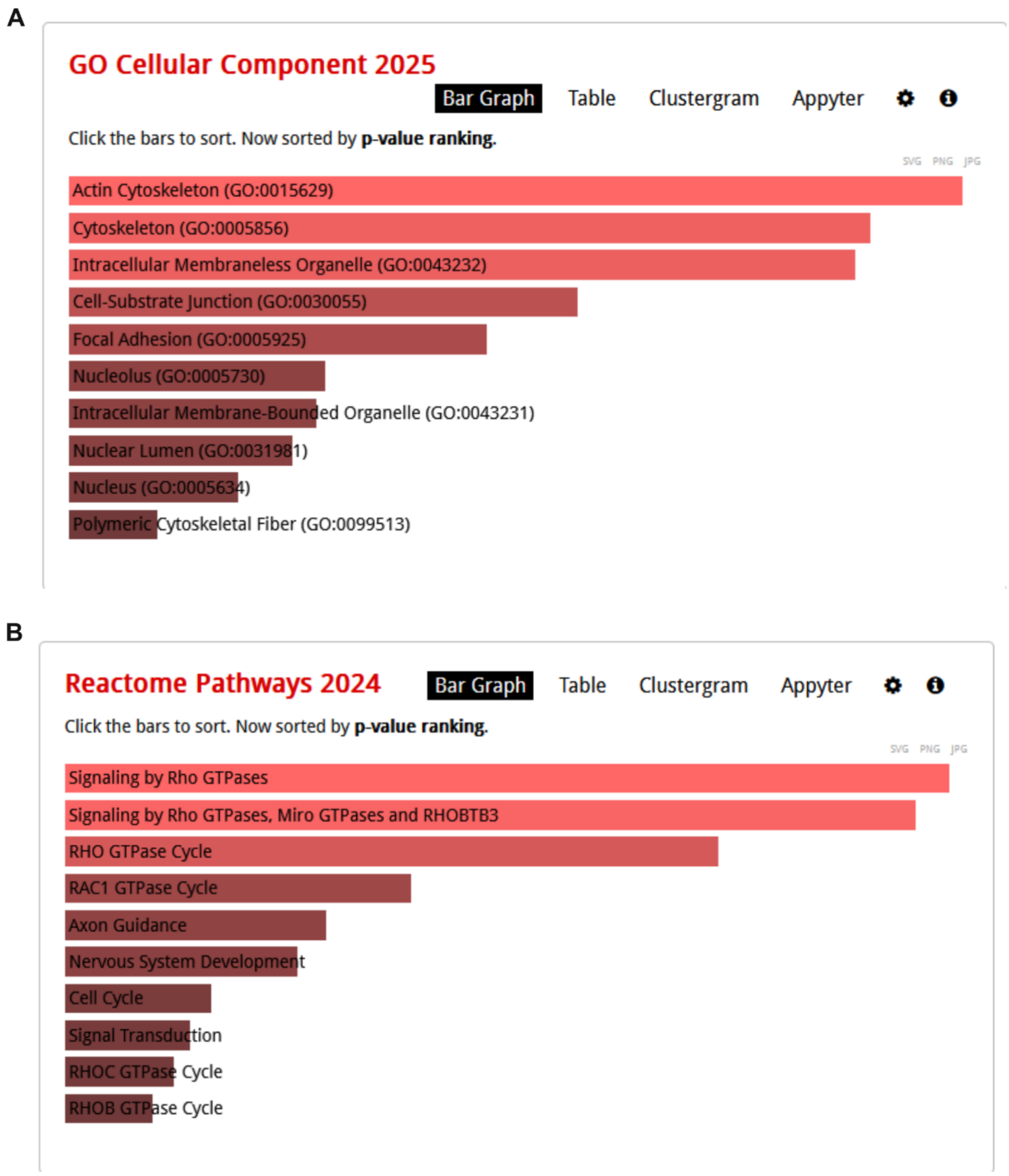


**Fig. S6. (A)** Gene Ontology and **(B)** Reactome Pathway analyses of predicted shared substrates of DMPK and LMTK3. Generated via Enrichr: https://maayanlab.cloud/Enrichr/enrich.

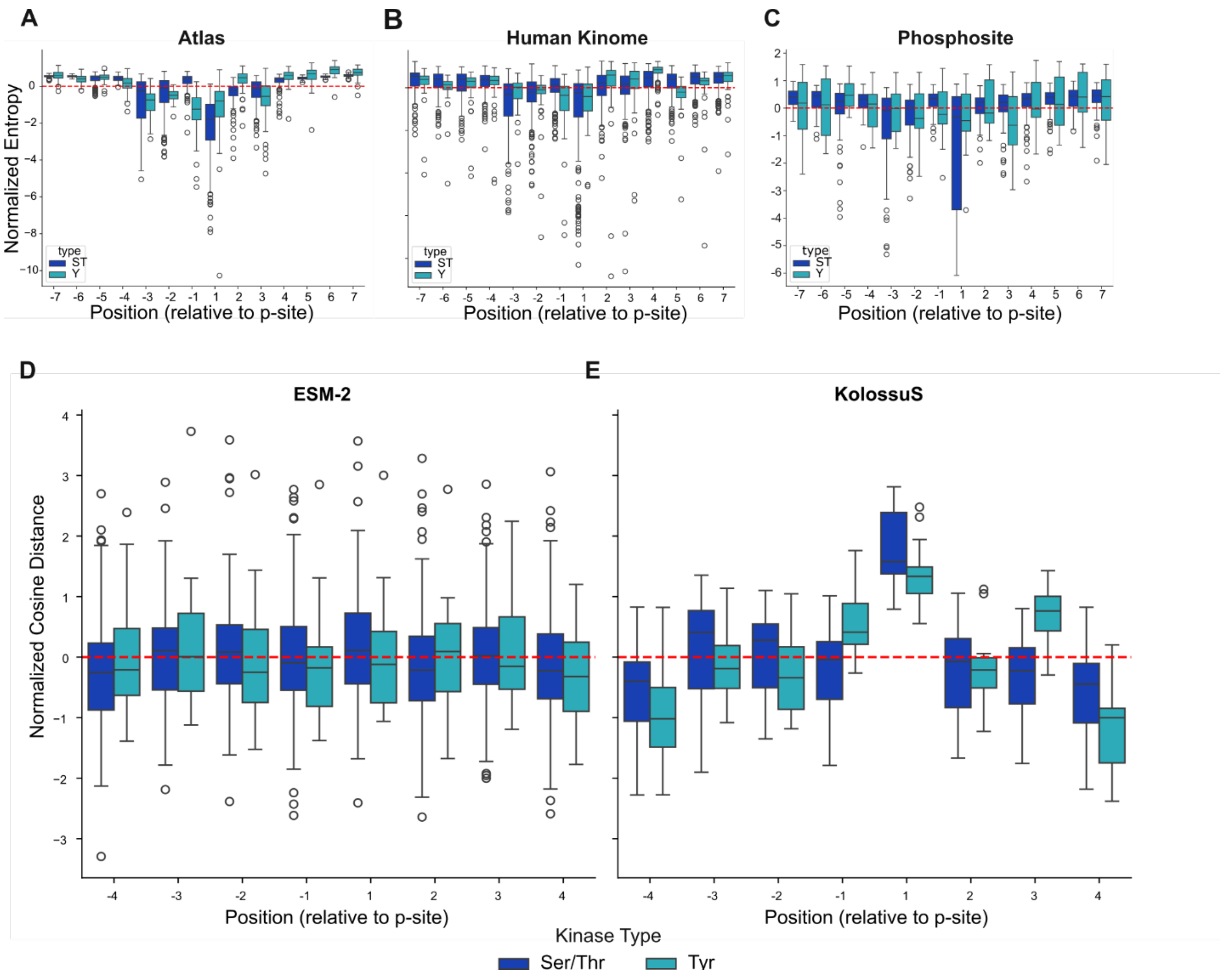


**Fig S7.** KolossuS embeddings capture the positional importance of substrate motif residues despite average pooling. **(A-C)** Position-specific entropy analysis on the Atlas, Human Kinome, and Phosphosite datasets, respectively, carried out separately for Ser/Thr (ST) and Tyr (Y) kinases (Materials and Methods). At positions –7 through +7, the boxplot indicates the distribution of normalized entropy scores observed at that position. A lower entropy score at a given position indicates greater preference towards specific amino acid residues at that position, while a higher entropy at a position indicates lower residue preference. The phosphorylation site (position 0) is omitted from the analysis, as by its very nature of only permitting S/T/Y residues it has low entropy. In Ser/Thr kinases, high residue preference is observed in the –3 and –2 positions, while Tyr kinases show high residue preference in the –1 and +3 positions. Both Ser/Thr and Tyr kinases exhibit high residue preference in the +1 position relative to phosphorylation. **(D-E)** In-silico perturbation analysis of the effect of substrate mutation in positions –4 to +4 relative to the phosphorylation site using ESM-2 and KolossuS embeddings (Materials and Methods). The cosine distance between mutated and wild-type embeddings was used to measure the expected change in kinase-binding specificity as a result of mutation in the substrate. In ESM-2, little positional effect of mutation is observed, while KolossuS shows a greater perturbation in substrate embeddings as a result of mutation in the –3 and –2 positions for Ser/Thr kinases, –1 and +3 for Tyr kinases, and +1 for both Ser/Thr and and Tyr kinases.

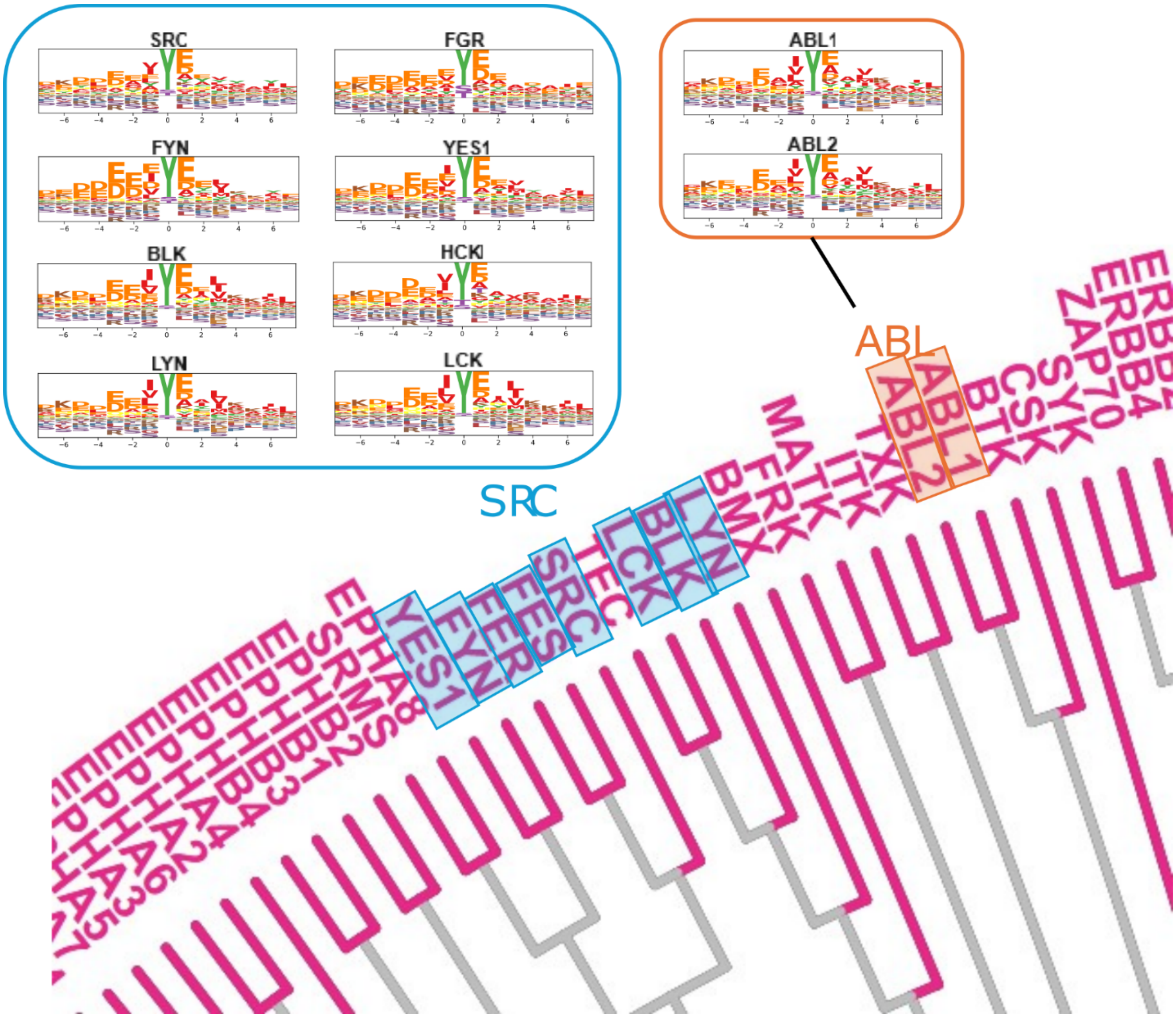


**Fig S8.** Validation of KolossuS-predicted kinase binding motifs for the SRC and ABL family of Tyrosine kinases. The SRC binding motifs demonstrate the preference for acidic residues in the –3 and –2, and +1 positions, and non-polar (valid, isoleucine) residues in the –1 and +3 positions of the substrate binding motif previously observed in (*4*) and (*5*). Similarly, the ABL binding motifs are consistent with the substrate specificity preferences reported in (*6*), and contains the sequence for “abltide”: “EAIYAAPFAKKK”, a BCR-ABL substrate frequently used as a positive control in assays involving substrate binding to the ABL kinase domain (*7*).

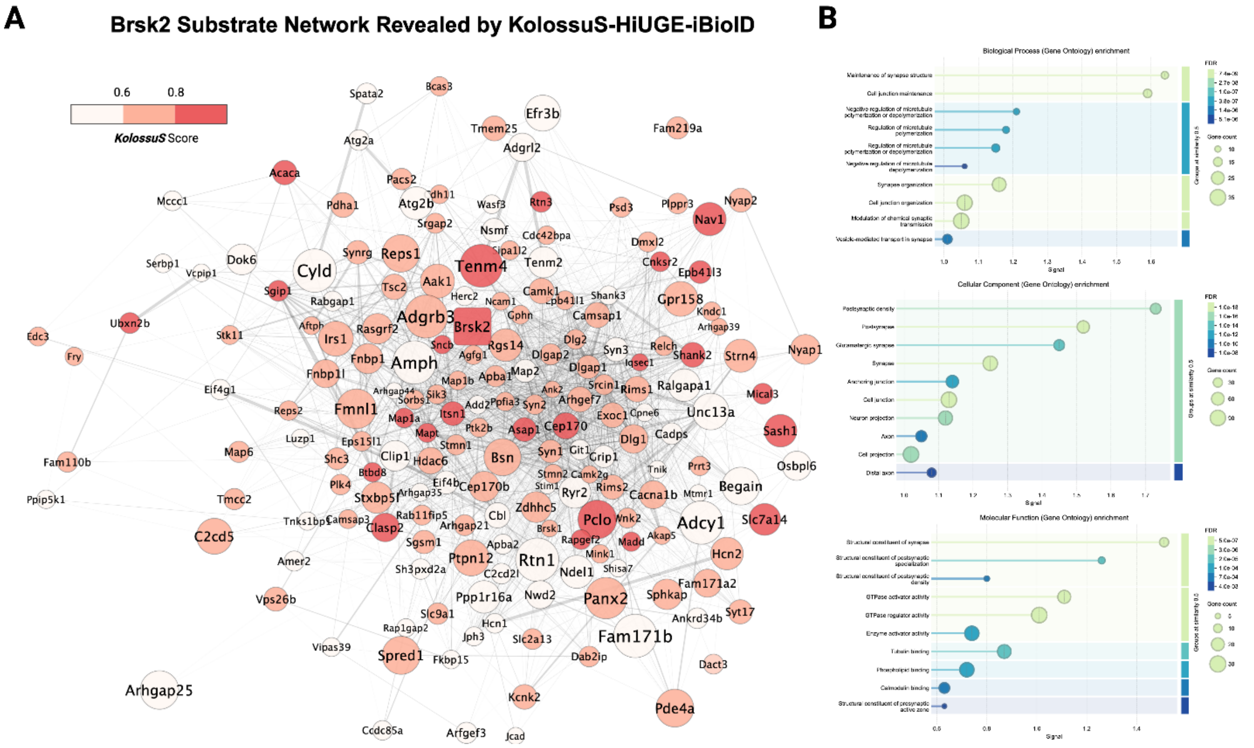


**Fig. S9. Brsk2 substrate network revealed by KolossuS-HiUGE-iBioID. (A)** Network representation of Brsk2 substrates identified through HiUGE-iBioID and filtered by KolossuS prediction scores (≥ 0.50). Node color indicates KolossuS scores, while node size reflects the number of unique phosphosites predicted for each protein. Edges represent protein-protein interactions obtained from the STRING database (medium confidence, 0.40). **(B)** Gene Ontology (GO) enrichment analysis of Brsk2 substrates (KolossuS ≥ 0.50), with enriched terms clustered by functional similarity using the Jaccard index via the STRING API (https://string-db.org/).

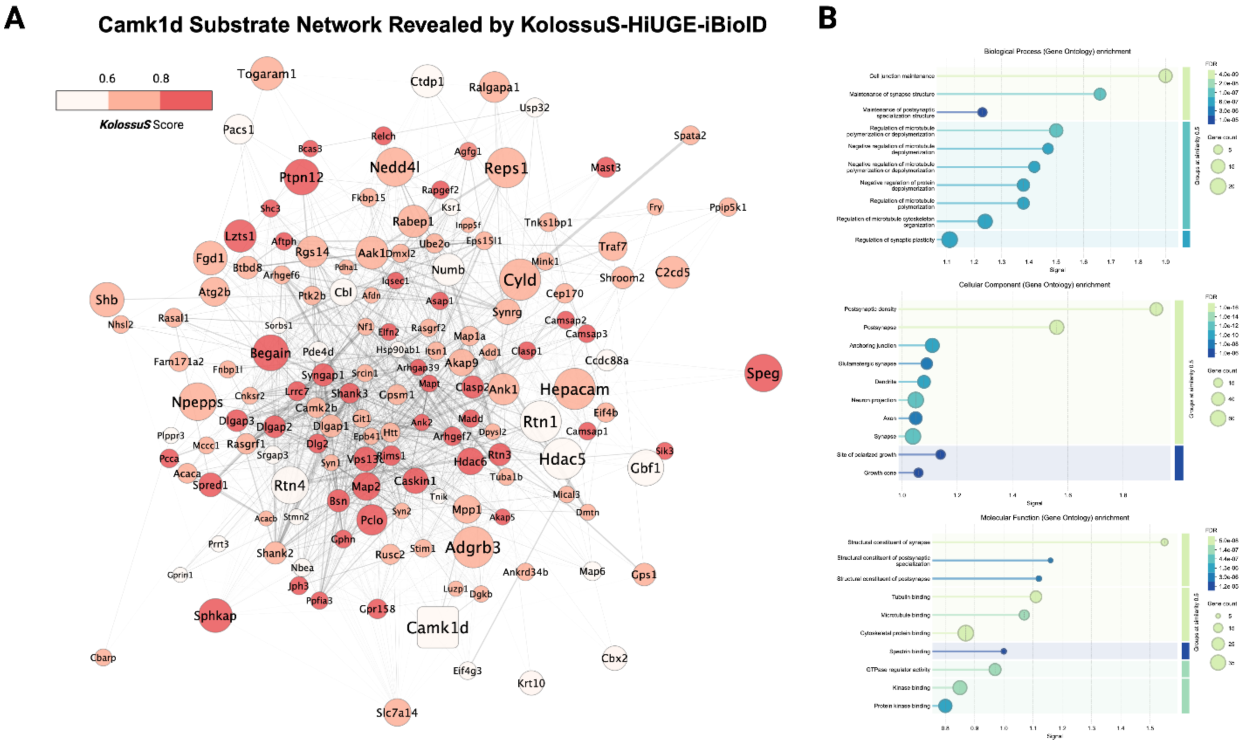


**Fig. S10. CaMK1D substrate network revealed by KolossuS-HiUGE-iBioID. (A)** Network representation of CaMK1D substrates identified through HiUGE-iBioID and filtered by KolossuS prediction scores (≥ 0.50). Node color indicates KolossuS scores, while node size reflects the number of unique phosphosites predicted for each protein. Edges represent protein-protein interactions obtained from the STRING database (medium confidence, 0.40). **(B)** Gene Ontology (GO) enrichment analysis of CaMK1Dsubstrates (KolossuS ≥ 0.50), with enriched terms clustered by functional similarity using the Jaccard index via the STRING API (https://string-db.org/).

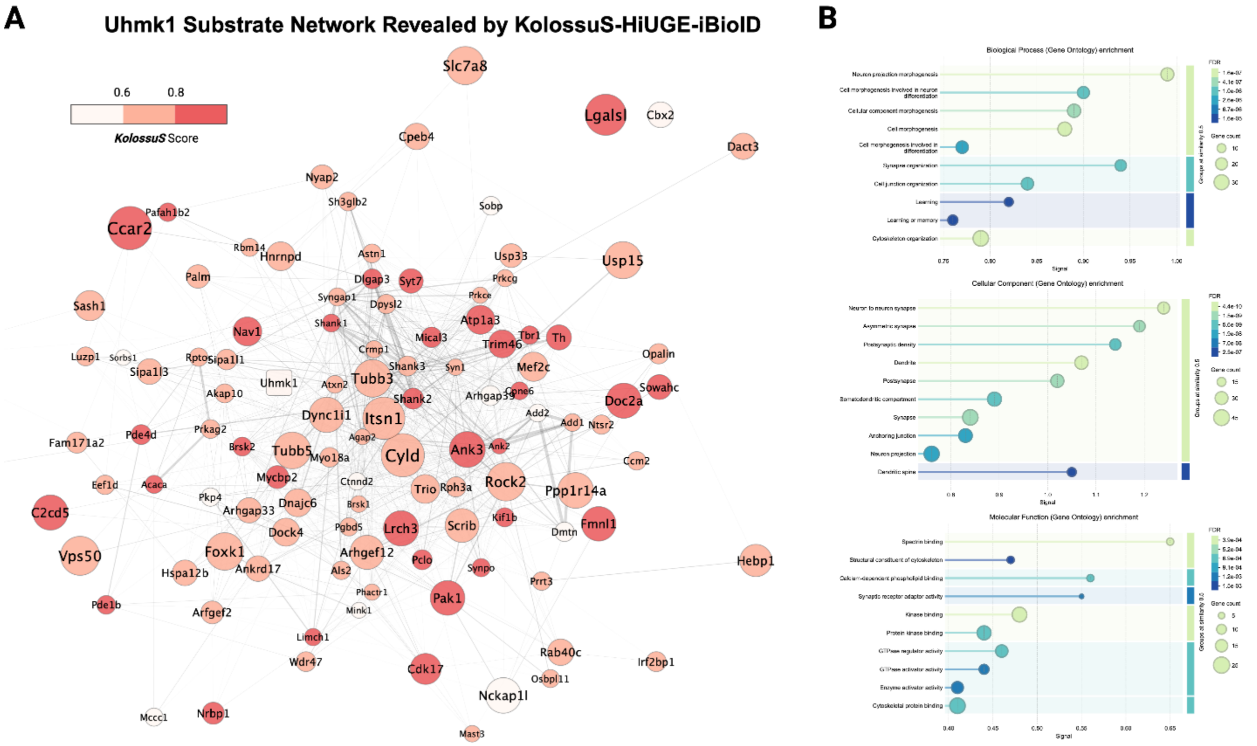


**Fig. S11. Uhmk1 substrate network revealed by KolossuS-HiUGE-iBioID. (A)** Network representation of Uhmk1 substrates identified through HiUGE-iBioID and filtered by KolossuS prediction scores (≥ 0.50). Node color indicates KolossuS scores, while node size reflects the number of unique phosphosites predicted for each protein. Edges represent protein-protein interactions obtained from the STRING database (medium confidence, 0.40). **(B)** Gene Ontology (GO) enrichment analysis of Uhmk1 substrates (KolossuS ≥ 0.50), with enriched terms clustered by functional similarity using the Jaccard index via the STRING API (https://string-db.org/).

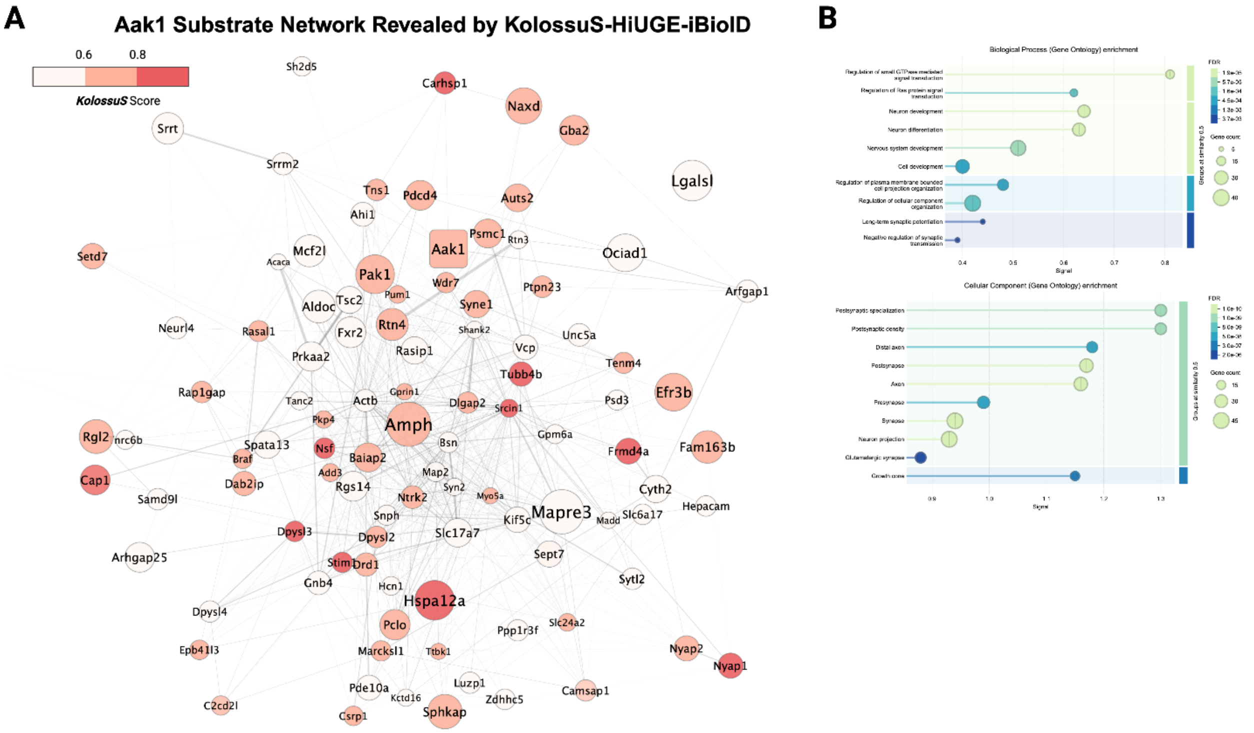


**Fig. S12. Aak1 substrate network revealed by KolossuS-HiUGE-iBioID. (A)** Network representation of Aak1 substrates identified through HiUGE-iBioID and filtered by KolossuS prediction scores (≥ 0.50). Node color indicates KolossuS scores, while node size reflects the number of unique phosphosites predicted for each protein. Edges represent protein-protein interactions obtained from the STRING database (medium confidence, 0.40). **(B)** Gene Ontology (GO) enrichment analysis of Aak1 substrates (KolossuS ≥ 0.50), with enriched terms clustered by functional similarity using the Jaccard index via the STRING API (https://string-db.org/).

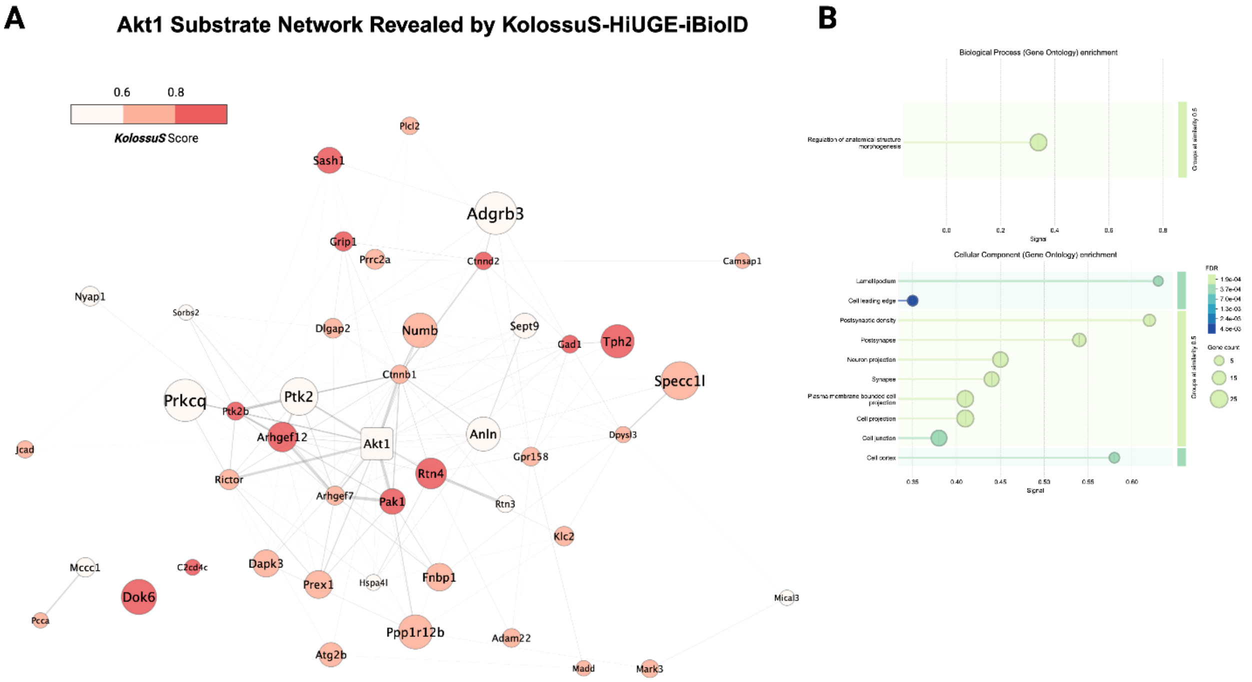


**Fig. S13. Akt1 substrate network revealed by KolossuS-HiUGE-iBioID. (A)** Network representation of Akt1 substrates identified through HiUGE-iBioID and filtered by KolossuS prediction scores (≥ 0.50). Node color indicates KolossuS scores, while node size reflects the number of unique phosphosites predicted for each protein. Edges represent protein-protein interactions obtained from the STRING database (medium confidence, 0.40). **(B)** Gene Ontology (GO) enrichment analysis of Akt1 substrates (KolossuS ≥ 0.50), with enriched terms clustered by functional similarity using the Jaccard index via the STRING API (https://string-db.org/).

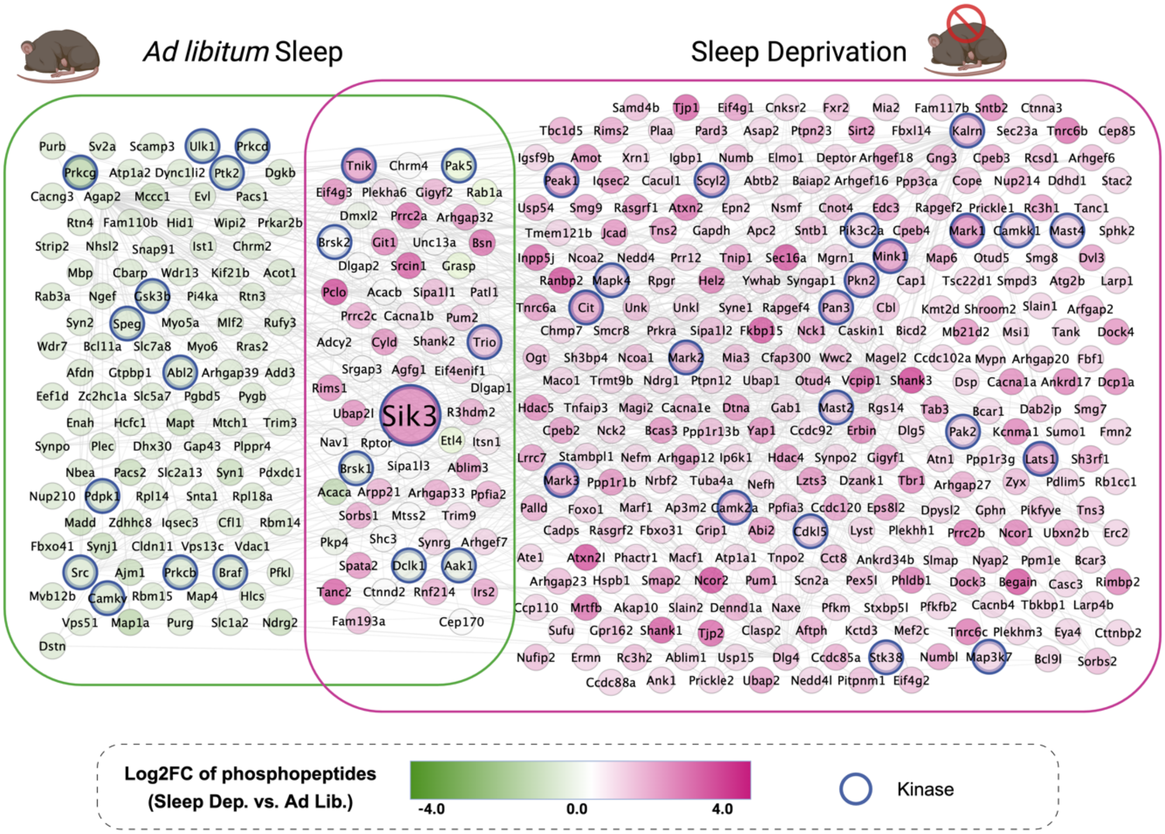


**Fig. S14. HiUGE-iBioID identification of Sik3 substrates under sleep deprivation.** Proximity phosphoproteome comparison of Sik3 under Ad libitum and sleep deprivation conditions. Proteins included as nodes were selected based on Log2FC > 1 and Bonferroni-adjusted p-value < 0.05. Node colors reflect the Log2 fold change in phosphorylated peptide abundance between sleep-deprived and Ad libitum samples, normalized to a protein-level ratio of one. Kinases other than Sik3 detected in proximity are highlighted with a blue border. Edges represent protein-protein interactions obtained from the STRING database (medium confidence, 0.40).

### Tables

| METRIC | Human | | | Mouse | | | Rat | | |
|---|---|---|---|---|---|---|---|---|---|
| | ESM-2 | Pre-FT | FT | ESM-2 | Pre-FT | FT | ESM-2 | Pre-FT | FT |
| Accuracy | 0.5640 | 0.6885 | 0.7117 | 0.4973 | 0.7265 | 0.7225 | 0.5096 | 0.6877 | 0.7145 |
| Sensitivity | 0.0586 | 0.7409 | 0.6189 | 0.0414 | 0.7982 | 0.6920 | 0.0489 | 0.8238 | 0.6990 |
| AUROC | 0.5026 | 0.7537 | 0.7617 | 0.4988 | 0.7905 | 0.7857 | 0.5183 | 0.7592 | 0.7741 |
| AUPR | 0.4254 | 0.6673 | 0.6792 | 0.4994 | 0.7562 | 0.7539 | 0.5039 | 0.7201 | 0.7389 |

**Table S1.** Performance on training ablation studies for human, mouse and rat Phosphosite data. ESM-2: ESM-2 15B embeddings; Pre-FT: KolossuS, trained only on the Atlas dataset without fine-tuning; FT: KolossuS, fine-tuned on the HumanKinome dataset.

| | Human | | Mouse | | Rat | |
|---|---|---|---|---|---|---|
| Metric | KolossuS | Phosformer-ST | KolossuS | Phosformer-ST | KolossuS | Phosformer-ST |
| Accuracy | 0.705356 | 0.65524 | 0.724932 | 0.677985 | 0.704136 | 0.637858 |
| Sensitivity | 0.63609 | 0.323724 | 0.725821 | 0.501369 | 0.68973 | 0.459459 |
| AUROC | 0.724932 | 0.749638 | 0.784734 | 0.783047 | 0.763463 | 0.735826 |
| AUPR | 0.725821 | 0.656128 | 0.755577 | 0.747761 | 0.728263 | 0.687475 |

**Table S2.** Performance benchmarks for KolossuS and Phosformer-ST on the Phosphosite evaluation data, provided for human, mouse, and rat Ser/Thr kinases.

| Family | F1-Max Threshold | | Max. F1-Score | |
|---|---|---|---|---|
| | **KolossuS** | **PhosformerST** | **KolossuS** | **PhosformerST** |
| AGC | **0.2542** | 0.0004 | 0.6644 | **0.6688** |
| Alpha | **0.2133** | 0.0002 | **0.6591** | 0.6289 |
| CAMK | **0.2957** | 0.0005 | 0.6420 | **0.6641** |
| CK1 | **0.2799** | 0.0001 | **0.6962** | 0.6521 |
| Other | **0.2878** | 0.0006 | **0.6665** | 0.6421 |
| PDHK | **0.3780** | 0.0007 | **0.5517** | 0.4400 |
| PIKK | **0.3571** | 0.0016 | **0.5748** | 0.5369 |
| STE | **0.2072** | 0.0002 | 0.6179 | **0.6220** |
| TKL | **0.1475** | 0.0002 | 0.6292 | **0.6333** |

**Table S3.** Score thresholds at which KolossuS and PhosformerST achieved their maximum F1-score (“F1-Max Threshold”). The F1-Score is a measure of classification accuracy that balances precision and recall. A perfectly calibrated model achieves its maximum F1-score at an F1-Max Threshold of 0.5. The maximum F1-Score achieved is also reported.

| | Aak1 | Akt1 | Brsk2 | Camk1d | Uhmk1 | Total |
|---|---|---|---|---|---|---|
| **Novel Peptides** | 186 | 57 | 190 | 116 | 79 | 628 |
| **Existing Peptides** | 953 | 154 | 833 | 525 | 192 | 2657 |
| **Total Peptides** | 1139 | 211 | 1023 | 641 | 271 | 3285 |
| **Total Proteins** | 674 | 182 | 400 | 284 | 205 | 1745 |

**Table S4.** Counts of phospho-peptides reported (Existing) or not reported (Novel) in PhosphoSitePlus for each of the HiUGE-iBioID-kinase phosphoproteomics of Aak1, Akt1, Brsk2, CaMK1D, and Uhmk1. Focused proximity phosphoproteomics results in deep coverage, including previously unreported phosphorylation sites.

| Gene | Site | Description | Feature Type |
|---|---|---|---|
| Ap3m2 | S287 | MHD | Domain |
| Arhgef6 | S487 | PH | |
| Elmo1 | S344 | ELMO | |
| Kcnma1 | S898 | RCK N-terminal 2 | |
| Nck2 | S351 | SH2 | |
| Pclo | S4470 | PDZ | |
| Rasgrf1 | S485 | PH 2 | |
| Rasgrf1 | S726 | N-terminal Ras-GEF | |
| Rasgrf2 | S484 | PH 2 | |
| Rasgrf2 | S745 | N-terminal Ras-GEF | |
| Shc3 | S410 | SH2 | |
| Tjp1 | S617/S686 | Guanylate kinase-like | |
| Arhgap32 | S1405/S1588 | Interaction with GAB2 | **Region involved in Interaction** |
| Arhgap32 | S1799 | Interaction with FYN | |
| Bsn | S2822/S2858/S3022/S3158 | Interaction with DAO | |
| Cbl | S667 | Interaction with CD2AP | |
| Cyld | S414 | Interaction with TRAF2 | |
| Cyld | S414/S556 | Interaction with TRIP | |
| Cyld | S556 | Interaction with IKBKG/NEMO | |
| Git1 | S419 | Interaction with NCK2 and GRIN3A | |
| Git1 | S419 | Interaction with PTK2/FAK1 | |
| Git1 | S601 | Interaction with IKBKG | |
| Hdac4 | S245 | Interaction with MEF2A | |
| Larp1 | S523 | Required for interaction with PABPC1 | |
| Nedd4 | S309 | Mediates interaction with TNIK | |
| Patl1 | S177 | Interaction with decapping machinery | |
| Sec16a | S1245/S1342/S1384 | Interaction with MIA3 | |
| Smap2 | S222 | Interaction with clathrin heavy chains | |
| Tnfaip3 | S553 | Interaction with TNIP1 | |
| Tnik | S649/S678 | Mediates interaction with NEDD4 | |
| Git1 | S419 | Required for localization at synapses | **Region involved in Localization** |
| Patl1 | S177 | Involved in nuclear foci localization | |
| Sec16a | S1245/S1342/S1384 | Required for endoplasmic reticulum localization | |

**Table S5. Structural features of phosphorylated sites targeted by Sik3.** List of phosphorylated residues identified as Sik3 substrates under sleep deprivation, as determined by KolossuS-HiUGE-iBioID. Each site is located within a protein domain or region previously associated with a specific function, as annotated in the UniProt database (https://www.uniprot.org/).
